# Developmental Recovery of Impaired Multisensory Processing in Autism and the Cost of Switching Sensory Modality

**DOI:** 10.1101/565333

**Authors:** Michael J. Crosse, John J. Foxe, Sophie Molholm

**Author notes:** Email addresses (M.J.C.), (J.J.F.), (S.M.).

## Abstract

Multisensory processing is often impaired in children with autism spectrum disorders (ASD) which may contribute to the social and communicative deficits that are prevalent in this population. Amelioration of multisensory deficits in adolescence has been observed for ecologically-relevant stimuli such as speech; however, this developmental recovery does not appear to generalize to the case of simple beeps and flashes, typically used in cognitive neuroscience research. Engagement of different neural processes and reduced environmental exposure to such nonsocial stimuli may lead to protracted multisensory development in neurotypical (NT) individuals, thus delaying the age at which individuals with ASD “catch up”. To determine whether recovery of multisensory deficits in ASD occurs later for ecologically-rare stimuli, we measured response times to randomly presented auditory, visual and audiovisual stimuli in over 350 participants between the ages of 6 to 40 years. By measuring the behavioral gain afforded by multisensory processing, we showed that individuals with ASD catch up to their NT peers by the mid-twenties. Computational modelling indicated that multisensory processing transitions from a default state of competition in early childhood, to one of facilitation in adulthood. This analysis also revealed developmental changes in sensory dominance that differed in individuals with ASD. Interestingly, we report that children and adolescents with ASD incurred less of a behavioral cost to switching sensory modality than their NT peers. This set of findings indicates that there is a complex interplay among the sensory systems that changes over the course of childhood, and differs in individuals with ASD.

## Introduction

Biological events tend to be multisensory, emanating or reflecting multiple forms of energy (e.g. photons, airborne vibrations, volatilized molecules, etc.). Humans have evolved a highly-specialized set of sensory receptors that enable us to sample these different forms of energy concurrently, optimizing how we perceive ecologically-relevant information. For instance, processing redundant multisensory signals often leads to faster reaction times (RTs) than processing the same information separately, a phenomenon known as the *redundant signals effect* (RSE; Todd, 1912; Hershenson, 1962; Kinchla, 1974). While a *race model* account of the RSE predicts that a response is triggered independently by the fastest sensory modality (Raab, 1962), the RSE typically exceeds the benefit predicted by mere statistical facilitation (Miller, 1982). Violation of the race model has been demonstrated using bisensory detection tasks for several decades and is widely interpreted as reflecting the multisensory gain due to pooled or integrated information processing (Gielen et al., 1983; Miller, 1986; Diederich and Colonius, 1987; Harrington and Peck, 1998; Molholm et al., 2002; Murray et al., 2004; Stevenson et al., 2012; Mégevand et al., 2013; Mahoney et al., 2015; Innes and Otto, 2019).

Whereas multisensory processing clearly influences how we perceive most biological events, particularly in instances when sensory evidence is ambiguous (Sumby and Pollack, 1954; Ross et al., 2007; Stevenson and James, 2009; Crosse et al., 2016), individuals with autism spectrum disorder (ASD) often do not benefit from the availability of multisensory information to the same extent as their neurotypical (NT) peers (de Gelder et al., 1991; Smith and Bennetto, 2007; Silverman et al., 2010; Irwin et al., 2011; Bebko et al., 2014; Stevenson et al., 2014a; Stevenson et al., 2014b; Foxe et al., 2015). We and others have suggested that impaired multisensory processing in ASD contributes to some of the commonly associated phenotypes such as atypical responses to sensory stimulation, and may even have detrimental effects on higher-order processes such as social interaction and communication (Ayres and Tickle, 1980; Martineau et al., 1992; Iarocci and McDonald, 2006; Foxe and Molholm, 2009; Beker et al., 2017; Stevenson et al., 2017).

In previous work by our lab, we demonstrated that multisensory gain increases steadily over the course of development using both simple audiovisual (AV) cues (Brandwein et al., 2011) as well as AV speech stimuli (Ross et al., 2011). Whereas multisensory processing was significantly impaired in children with ASD for both of these tasks (Brandwein et al., 2013; Foxe et al., 2015), NT levels of AV speech integration are achieved by the time individuals with ASD reach adolescence (Taylor et al., 2010; Foxe et al., 2015). In contrast, high-functioning teenagers with ASD failed to show reliable multisensory gain when performing a simple AV detection task (Brandwein et al., 2013). Recent theoretical (Beker et al., 2017) and computational (Cuppini et al., 2017) perspectives have suggested that the constant exposure to AV speech during maturation may serve to train multisensory speech function, leading to earlier developmental recovery of function in ASD. Furthermore, the underlying neural processes that are engaged in different contexts may follow disparate developmental trajectories; the trajectory of multisensory development in NT individuals reaches full maturity much earlier for speech stimuli (Ross et al., 2011) compared to non-speech stimuli (Brandwein et al., 2011). Here, using the same bisensory detection task, we tested the hypothesis that recovery of multisensory function in ASD occurs at a later developmental stage for nonsocial stimuli.

Electrophysiological studies in the cat superior colliculus (SC) has shown that the ability of SC neurons to integrate multisensory inputs cooperatively is not present at birth (Wallace and Stein, 1997, 2001), but rather emerges and matures in the immature nervous system with exposure to multisensory experiences (Wallace et al., 2004; Wallace and Stein, 2007; Yu et al., 2010; Xu et al., 2014). Computational modelling suggests that multisensory signals interact by default in a competitive manner, inhibiting effective processing of redundant stimuli (Yu et al., 2019). Considerable postnatal exposure to multisensory cues is thought to strengthen excitatory cross-sensory projections, promoting interactions of a facilitative nature (Cuppini et al., 2011; Cuppini et al., 2012; Cuppini et al., 2018). While numerous developmental studies in humans have reported reduced multisensory ability in young children (Gori et al., 2008; Barutchu et al., 2009), there is little empirical evidence of such competitive multisensory interactions other than that reported in adults (Molholm et al., 2004; Sinnett et al., 2008). Here, we used a computational approach to directly test whether multisensory behavior in children reflected competitive or facilitative interactions. The type of interaction was determined by how accurately hypothesis-driven models of multisensory processing could predict empirical multisensory benefits (Otto et al., 2013; Innes and Otto, 2019). The same modelling approach was used to quantify developmental changes in sensory dominance and any potential group differences therein. We expected that such inherent sensory weighting would have a greater impact on interactions of a competitive nature.

When switching from one sensory modality to another, average response times are slower on trials preceded by a different sensory modality (switch trials) compared to trials preceded by the same modality (repeat trials; Wundt, 1893; Sutton et al., 1961; Spence et al., 2001). Modality switch effects (MSEs) are inherent to any bisensory detection task that uses an intermixed stimulus presentation design (Gondan and Minakata, 2016; Otto and Mamassian, 2017) and have been shown to systematically contribute to the RSE because they are typically larger on unisensory trials than on multisensory trials (Gondan et al., 2004; Van der Stoep et al., 2015a; Shaw et al., 2019). Moreover, data suggest that children with high-functioning ASD incur a greater cost when switching from auditory to visual stimuli than their NT peers (Williams et al., 2013). We therefore considered group differences in MSEs and quantified their contribution to the RSE. Using a computational modelling framework (Otto and Mamassian, 2012), we measured the dependency between RTs on different sensory channels, giving us insight into how attention was spread across sensory inputs during speeded bisensory detection, and considered how this in turn could impact MSEs. We discuss the implications of MSEs on the interpretation of the RSE, and how the interplay between multisensory integration and switch effects may contribute differentially over the course of development in NT and ASD individuals.

## Methods

The present study is based on new analyses of a large body of data collected as part of several previously published studies (Brandwein et al., 2011; Brandwein et al., 2013; Brandwein et al., 2015), as well as new unpublished data.

### Participants

A total of 400 individuals participated in the experiment. The data of 42 participants (10.5% of the total sample, 29 ASD) were excluded from all analyses based on the following criteria: 1) they did not fall within the desired age range of 6–40 years, 2) their performance IQ was below 80, 3) their detection accuracy was less than 3 SDs below the sample’s mean, 4) they had an excessive number of false alarms, 5) they had a disproportionate number of misses on visual trials (excessive eye-closure) or on audio trials (not listening), or 6) they had less than 20 RTs per condition (this can bias median RT estimates (Miller, 1988, 1991) as well as race model estimates (Kiesel et al., 2007)). Of the remaining 358 participants, 225 met criteria for NT (age range: 6–36 years; 115 females) and 133 had a diagnosis of ASD (age range: 6–39 years; 34 females). For analysis purposes, age was either treated as a continuous variable or participants were cross-sectioned into four developmental subgroups: children (6–9 years), pre-adolescents (10–12 years), adolescents (13–17 years), adults (18–40 years). Mean age was not statistically different between NT and ASD participants in any of the four age groups (*t* < 0.91, *p* > 0.38, *d* < 0.21). Participant demographics are presented in Table 1.

**Table 1.** Demographic characteristics of participant populations.

|  | NT |  |  |  | ASD |  |  |  |
| --- | --- | --- | --- | --- | --- | --- | --- | --- |
|  | 6–9 yrs | 10–12 yrs | 13–17 yrs | 18–40 yrs | 6–9 yrs | 10–12 yrs | 13–17 yrs | 18–40 yrs |
| *n* | 51 | 46 | 54 | 74 | 44 | 34 | 31 | 23 |
| *n*<sub>female</sub> | 27 | 26 | 24 | 38 | 7 | 6 | 10 | 11 |
| *n*<sub>IQ</sub> | 45 | 43 | 48 | 10 | 44 | 33 | 30 | 21 |
| Age | 8.1 (1.2) | 11.5 (1.0) | 15.0 (1.3) | 25.3 (3.6) | 8.1 (1.0) | 11.4 (0.7) | 14.7 (1.6) | 25.8 (5.1) |
| F<sub>1</sub> score | 0.90 (0.07) | 0.93 (0.06) | 0.95 (0.04) | 0.97 (0.02) | 0.85 (0.08) | 0.87 (0.08) | 0.92 (0.07) | 0.95 (0.04) |
| PIQ | 106.1 (13.0) | 109.7 (10.7) | 104.9 (13.3) | 109.9 (12.3) | 106.2 (17.1) | 106.6 (16.2) | 107.9 (13.2) | 108.5 (13.9) |
| VIQ | 113.0 (10.6) | 111.8 (13.0) | 113.1 (12.8) | 115.1 (16.1) | 97.3 (19.8) | 99.4 (19.1) | 99.9 (18.8) | 109.3 (15.8) |
| FSIQ | 111.4 (11.5) | 112.2 (11.7) | 110.1 (12.5) | 114.5 (14.0) | 101.7 (17.5) | 102.8 (17.4) | 104.1 (14.1) | 110.0 (14.4) |
| ADOS | – | – | – | – | 7.3 (2.3) | 8.0 (0.9) | 6.9 (3.3) | – |

Note: $n_{\text{female}}$ indicates the number of female participants in respective age groups and $n_{\text{IQ}}$ indicates the number of participants for whom IQ scores were obtained. The number of participants for whom ADOS scores were obtained is 31, 19, 7 respectively. PIQ: performance IQ; VIQ: verbal IQ; FSIQ: full-scale IQ (assessed using the WASI). $F_1$ scores indicate participants' detection accuracy, accounting for false alarms (see Methods for details). Values indicate the group mean with standard deviation shown in parentheses.

Individuals were excluded from participating in the experiment if they had a history of seizures or head trauma, or a known genetic disorder. Additionally, NT participants were excluded if they had a history of psychiatric, educational, attentional or other developmental difficulties (as assessed by a history questionnaire), a biological first-degree relative with a known developmental disorder, or if they or their legal guardians endorsed six or more items of inattention or hyperactivity on a DSM-IV checklist for attention deficit disorder. For the vast majority of participants, diagnoses of ASD were obtained by a trained clinical psychologist using the Autism Diagnostic Interview-Revised (Lord et al., 1994) and the Autism Diagnostic Observation Schedule (ADOS; Lord et al., 2000). Diagnoses of the remaining individuals were made by a licensed clinical psychologist external to this study using the Diagnostic Criteria for Autistic Disorder from the DSM-IV TR (APA, 2000). For more details regarding sub-phenotyping, medication and ethnic demographics, please refer to Brandwein et al. (2013) and Brandwein et al. (2015).

IQ quotients for performance (PIQ), verbal (VIQ) and full-scale (FSIQ) intelligence were assessed in the majority of participants using the Wechsler Abbreviated Scales of Intelligence (WASI; Stano, 1999). Note that mean PIQ was not statistically different between NT and ASD participants in any of the four age groups (*t* < 1.2, *p* > 0.23, *d* < 0.27). The descriptive statistics for each of the subgroups are summarized in Table 1. Participants were formally screened for normal or corrected-to-normal vision using a Snellen eye test chart and audiometric threshold evaluation confirmed that all participants had within-normal-limits hearing. All procedures were approved by the institutional review boards of the City College of New York and the Albert Einstein College of Medicine. All participants or legal guardians of participants provided written informed consent in accordance with the tenets of the 1964 Declaration of Helsinki.

### Stimuli and procedure

The stimulus materials were identical to those described in Brandwein et al. (2011). In brief, visual (V) stimuli consisted of a red disc (diameter: 3.2 cm; duration: 60 ms), located 0.4 cm above a central fixation crosshair on a black background. The disc subtended visual angles of 1.5° vertically and horizontally and the bottom of the disc subtended 0.9° vertically above the crosshair (Fig. 1A). Auditory (A) stimuli consisted of a 1-kHz pure tone, sampled at 44.1 kHz (duration: 60 ms; rise/fall time: 5 ms). Audiovisual (AV) stimuli consisted of the combined simultaneous pairing of the auditory and visual stimuli described above.

**Figure 1.**
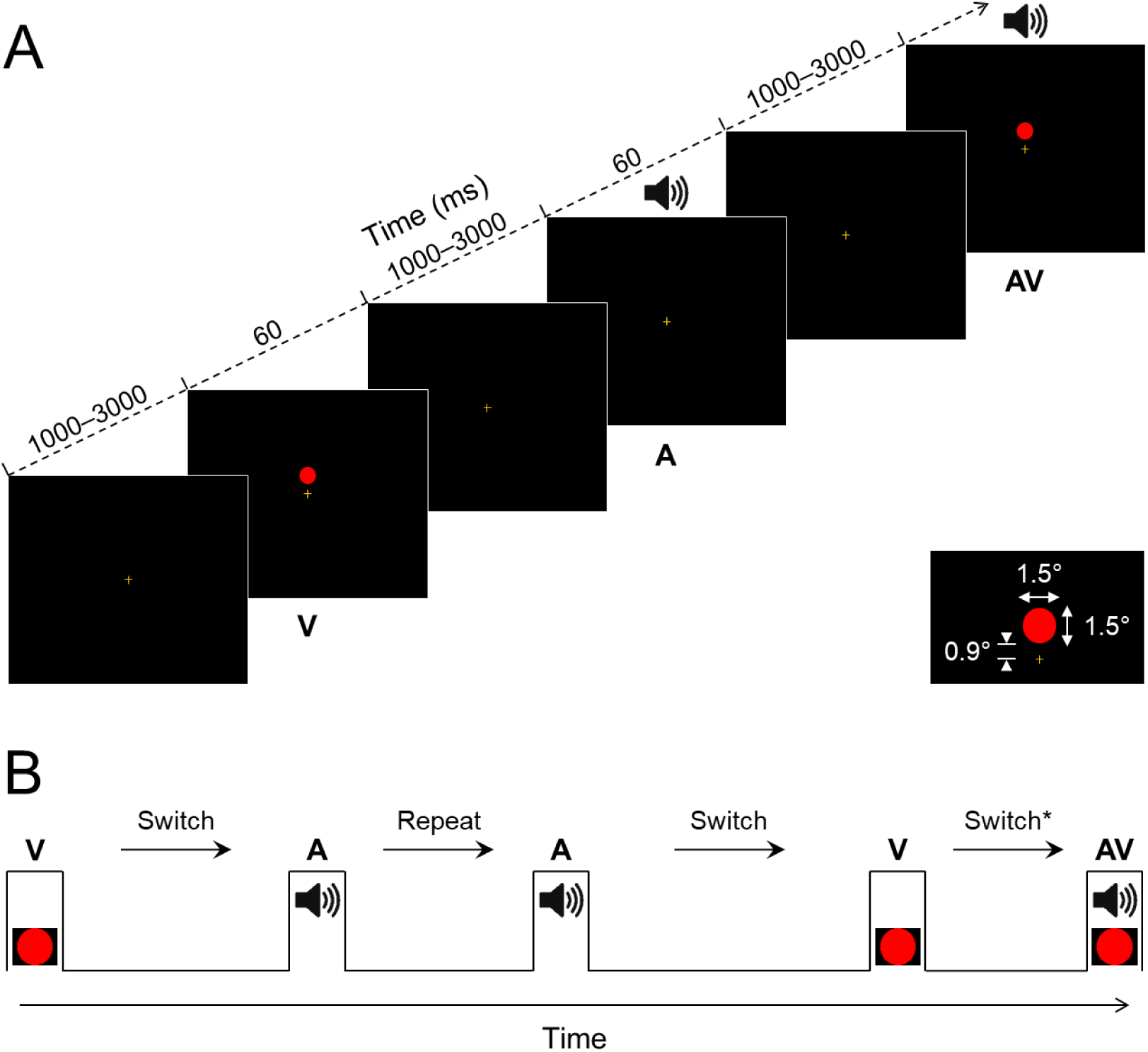
Bisensory detection task. **A**, Auditory (A), visual (V) and audiovisual (AV) stimuli (60-ms duration) were presented in a randomized order every 1000–3000 ms. Participants responded to each stimulus with a button press as fast as possible. **B**, Stimuli were categorized as either switch or repeat trials based on the modality of the preceding stimulus (repeat trials: AV→AV, A→A, V→V; switch trials: V→AV*, A→AV*, A→V, V→A). Asterisks indicate trials that are only partial switches. Trials AV→A and AV→V were excluded from the analysis as they were considered neither switches nor repeats.

Participants performed a speeded bisensory detection task on a computer and were seated 122 cm from the visual display in a dimly-lit, sound-attenuated booth. RTs were recorded during the simultaneous recording of electrophysiological (EEG) data, however, the EEG data are not reported in this study (for an account of previous EEG analyses, please refer to Brandwein et al., 2011; Brandwein et al., 2013; Brandwein et al., 2015). To reduce predictability, the stimuli were presented in a completely randomized order with equal probability and the interstimulus interval (ISI) was randomly jittered between 1000–3000 ms according to a uniform, square-wave distribution (see Fig. 1A). Stimulus presentation was controlled using Presentation® software (Neurobehavioral Systems, Inc., Berkeley, CA). Auditory stimuli were delivered binaurally at an intensity of 75 dB SPL via a single, centrally-located loudspeaker (JBL Duet Speaker System, Harman Multimedia). Visual stimuli were presented at a resolution of 1280 × 1024 pixels on a 17-inch Flat Panel LCD monitor (Dell Ultrasharp 1704FTP). The auditory and visual stimuli were presented in close spatial proximity, with the speaker placed atop the monitor and aligned vertically to the visual stimulus. Participants were instructed to press a button on a response pad (Logitech Wingman Precision Gamepad) with their right thumb as soon as they perceived any of the three stimuli. Analogue triggers indicating the latencies of stimulus onsets and button presses were sent to the acquisition PC via Presentation® and stored digitally at a sampling rate of 512 Hz in a separate channel of the EEG data file using ActiView software (BioSemi^TM^, Amsterdam, The Netherlands). Stimuli were presented in blocks of ∼100 trials and participants typically completed 6–10 blocks in total.

### Data analysis

Detection accuracy was assessed in order to identify participants that did not attend adequately to the stimuli. To account for false alarms and excessive button pressing, F_1_ scores were computed as the harmonic mean of precision and recall (Van Rijsbergen, 1979):

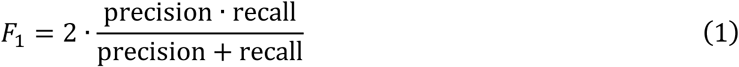

where precision = hits/(hits + false alarms) and recall = hits/(hits + misses). Responses were considered as false alarms if they occurred earlier than 100 ms post stimulus onset, or if they occurred after another response but before the next stimulus. Responses were considered as misses if they occurred later than 2000 ms post stimulus onset, or if there was no response at all to a given stimulus.

Response times were measured relative to the onset time of the preceding stimulus and analyzed separately for each participant in MATLAB (The MathWorks, Inc., Natick, MA). Responses were excluded from all analyses if there was more than one response within a given trial (double-presses), they occurred within the first 3 trials of a block (considered training) or the preceding ISI was not between 1000–3000 ms (due to system errors). An outlier correction procedure was performed before the main RT analyses. First, RTs that did not fall within 100–2000 ms post-stimulus were removed. On average, fast outliers (<100 ms, considered anticipatory responses) made up 0.7% (±0.9) of trials and slow outliers (>2000 ms, considered misses) made up 0.4% (±0.6) of trials. Second, RTs outside the middle 95^th^ percentile (2.5– 97.5) of their respective conditions were removed. While not all RTs outside of this range are necessarily outliers, those within this range are most likely to come from the cognitive processes under consideration (Ratcliff, 1993). This approach minimizes the impact of outliers with only negligible truncation biases (Ulrich and Miller, 1994) and captures the range of RTs at an individual-participant level, an important factor when dealing with significant inter-subject variability.

Analysis of RT data was conducted on the whole RT distribution by splitting it into discrete quantiles (Ratcliff, 1979). RTs were organized into 20 linearly-spaced quantiles between the 2.5–97.5 cutoffs used for outlier correction. Because outlier correction was performed separately for each condition, the lowest 2.5 and highest 97.5 percentiles were used for all three conditions in order to maintain the relationship between them. Cumulative distribution functions (CDFs) were obtained by calculating the cumulative probability of RTs occurring below time *t* given a signal X, *P*(RT_X_ ≤ *t*|X). CDFs were averaged or “Vincentized” across participants at each corresponding quantile (Vincent, 1912). Note, this approach does not require there to be an equal number of RTs in each condition (Ulrich et al., 2007).

### Race model analysis

To obtain quantitative predictions of statistical facilitation, we used Raab’s race model (Raab, 1962). Race models predict that the response to a redundant signal is triggered independently by the fastest sensory modality. Let *P*(RT_A_ ≤ *t*|AV) and *P*(RT_V_ ≤ *t*|AV) represent the CDFs of the A and V components of an AV stimulus, respectively. Assuming the RT distributions of the A and V components overlap, the probability of either triggering a response can be represented using probability summation:

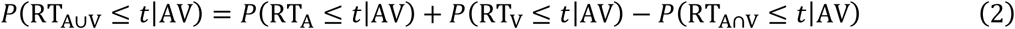

where *P*(RT_A∩V_ ≤ *t*|AV) is the probability of the A and V signals triggering a response at the same time. To solve this analytically, we need to make two assumptions: 1) RTs to the A and V components of the AV signal follow the same distributions as the RTs to the unisensory A and V signals, such that *P*(RT_A_ ≤ *t*|AV) = *P*(RT_A_ ≤ *t*|A) and *P*(RT_V_ ≤ *t*|AV) = *P*(RT_V_ ≤ *t*|V), an assumption known as context invariance (Ashby and Townsend, 1986; Luce, 1986; Miller, 2016); 2) RTs to the A and V components of the AV signal are statistically independent, such that their joint probability *P*(RT_A∩V_ ≤ *t*|AV) can be calculated by the product of *P*(RT_A_ ≤ *t*|AV) and *P*(RT_V_ ≤ *t*|AV) (Meijers and Eijkman, 1977). Simplifying *P*(RT_A∪V_ ≤ *t*|AV) to *F*_A∪V_(*t*), *P*(RT_A_≤ *t*|A) to *F*_A_(*t*) and *P*(RT_V_ ≤ *t*|V) to *F*_V_(*t*), equation 2 can be represented as:

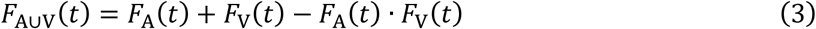

Note, the joint probability term is often omitted from Equation 3 to produce an upper bound known as Miller’s bound or the race model inequality (Miller, 1982), as the assumption of statistical independence is poorly motivated; it is likely that responses to signals on different sensory channels compete for resources (Miller, 1978, 1982; Colonius, 1986, 1990; Gondan and Minakata, 2016). Assuming that the allocation of attentional resources to each channel is partially determined by the modality of the previous trial (Miller, 1982), we separated the unisensory RTs by their preceding sensory modality and computed individual race models before taking their average:

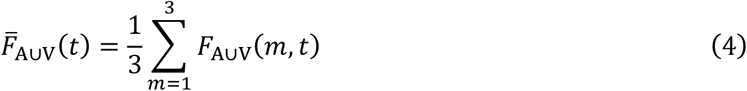

where *m* is the preceding modality. This approach captured some of the dependency between RTs to signals on different channels, resulting in an estimate of statistical facilitation that was less conservative at every quantile (*p* < 0.025, two-tailed permutation tests). Note that using Raab’s model or Miller’s bound typically yields the same outcome qualitatively (Van der Stoep et al., 2015b; Van der Stoep et al., 2015a).

Multisensory benefits were quantified by the area between the CDFs in the multisensory condition and the most effective unisensory condition (Otto et al., 2013). First, we computed the multisensory benefit predicted by the race model (Fig. 2B, left):

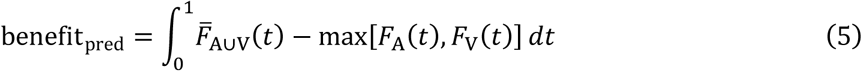

where the integral is taken over every quantile *t* from 0 to 1. The term max[*F*_A_(*t*), *F*_V_(*t*)] represents a lower bound of facilitation, known as Grice’s bound (Grice et al., 1984), whereby no statistical benefit is observed for a redundant signal at any quantile. Similarly, we computed empirical benefits based on the actual multisensory RTs (Fig. 2B, right):

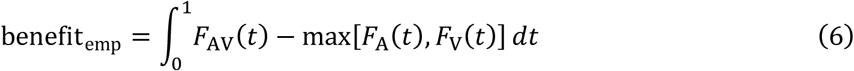

Note that this is not the same as measuring multisensory interactions since Grice’s bound does not account for statistical facilitation (see Innes and Otto, 2019). Rather, it quantifies the benefit afforded by a redundant signal relative to that of the most effective unisensory signal.

**Figure 2.**
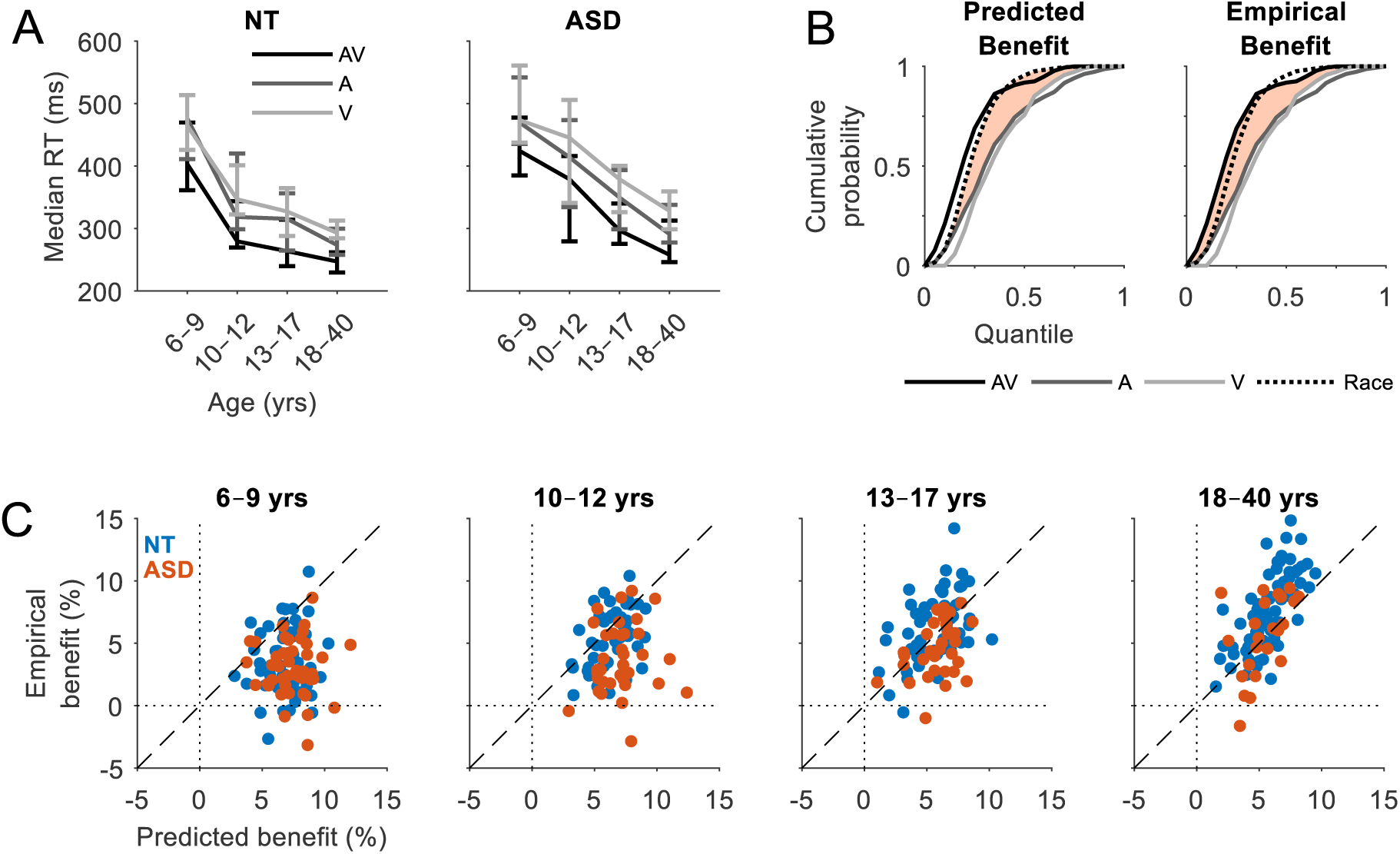
Reaction times and multisensory benefits. **A**, Group median RTs for NT (left panel) and ASD (right panel) individuals as a function of age group. Error bars indicate 95% CIs (bootstrapped). **B**, RT cumulative probability for each of the three stimulus conditions and the race model (Eq. 4). Predicted benefits (left panel) are quantified by the area between the CDFs of the race model and the faster of the unisensory conditions (Eq. 5). Empirical benefits (right panel) are quantified by the area between the CDFs of the multisensory condition and the faster of the unisensory conditions (Eq. 6). Data from an example NT adult participant. **C**, Predicted benefits versus empirical benefits by age group. Each datapoint represents an individual participant (blue = NT, red = ASD).

**Figure 3.**
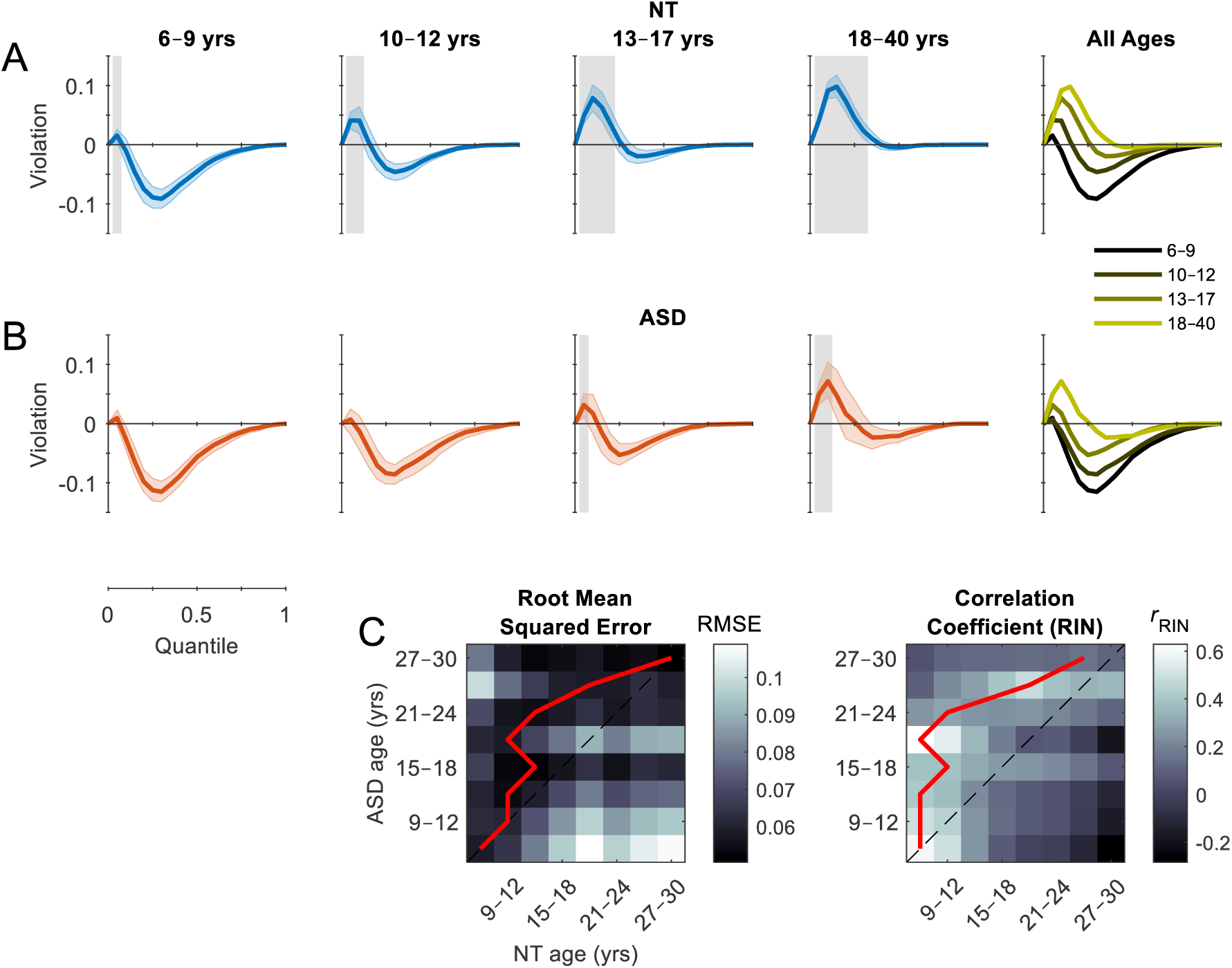
Testing the race model. **A**, **B**, Violation of the race model is quantified by the difference between the CDFs of the multisensory condition and the race model. Positive values reflect quantiles where multisensory RTs were faster than predicted by the race model. Gray shaded regions indicate significant differences (*p* < 0.05, right-tailed permutation tests, *t*_max_ corrected). Colored error bounds indicate 95% CIs (bootstrapped). **C**, Root mean squared error (left panel) and RIN-transformed Pearson correlation coefficient (right panel) between the violation functions for NT and ASD participants of different ages (range: 6–30 years, increment: 3 years). Red lines indicate the minimum (left panel) and maximum (right panel) values of each row (i.e., the age groups that were most similar). Divergence of the red line above the dotted midline indicates a developmental delay in ASD participants.

To determine whether the empirical multisensory benefits exceeded statistical facilitation, we computed the difference between the CDFs of the multisensory condition and the race model at every quantile (Molholm et al., 2002; Molholm et al., 2006). Positive values indicate quantiles where multisensory RTs were faster than predicted, i.e., violation of the race model. To obtain an overall index of multisensory gain, we calculated the area under the curve (AUC) by taking the integral over every quantile as before (Fig. 4A):

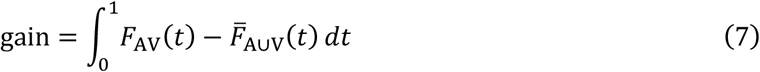

**Figure 4.**
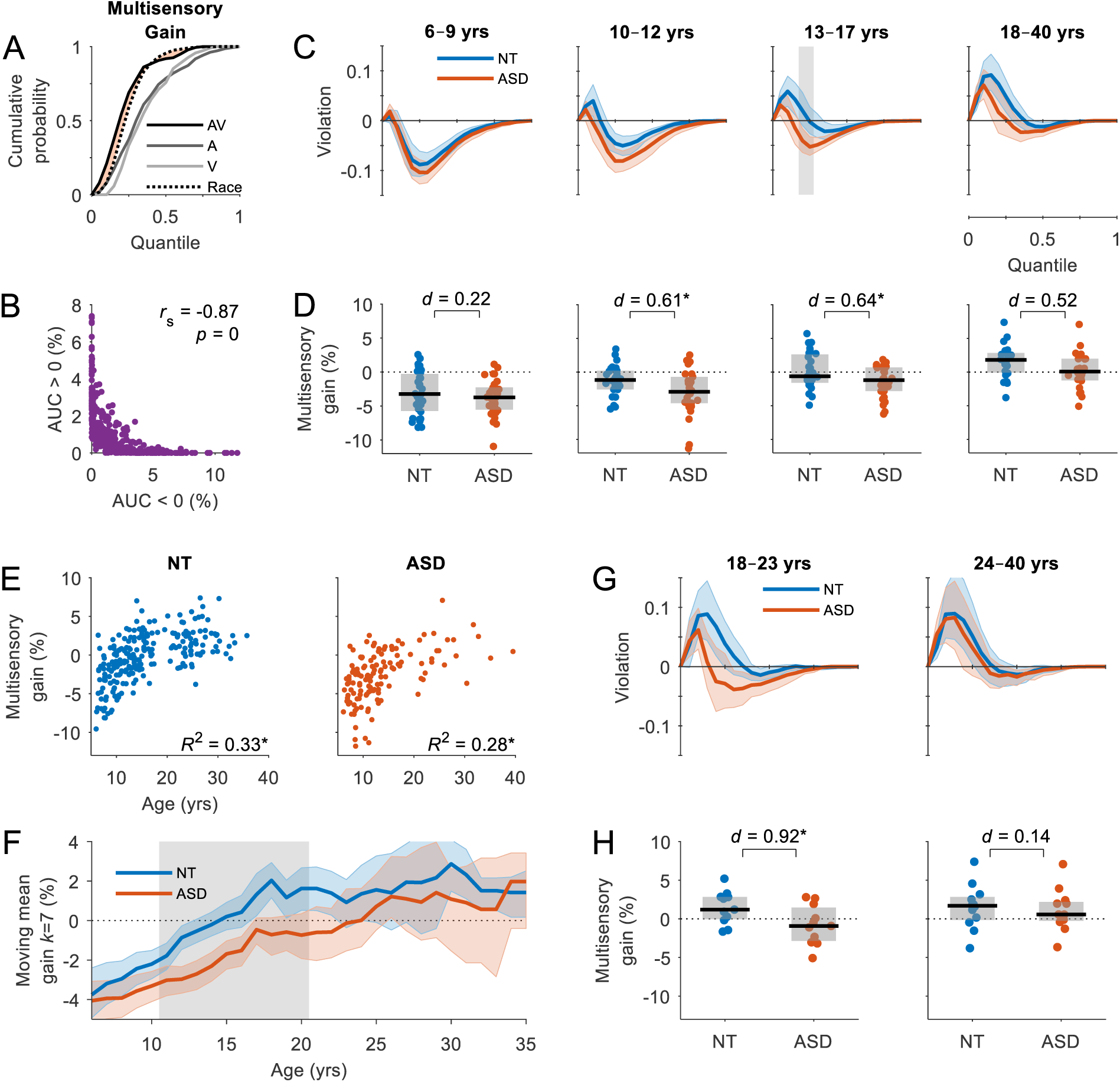
Developmental course of multisensory gain. **A**, RT cumulative probability for each of the three stimulus conditions and the race model. Multisensory gain is quantified by the area between the CDFs of the multisensory condition and the race model (Eq. 7). Data from an example NT adult participant. **B**, The area under the curve (AUC) below zero is negatively correlated with the AUC above zero, providing information about participants that do not exceed statistical facilitation. **C**, Race model violation for ASD (red trace) and sex-and age-matched NT (blue trace) participants by age group. Colored error bounds indicate 95% CIs (bootstrapped). Gray shaded regions indicate significant group differences (*p* < 0.05, two-tailed permutation tests, *t*_max_ corrected). **D**, Multisensory gain by age group. Boxplots indicate the median value (black line) and interquartile range (grey box). Each datapoint represents an individual participant (blue = NT, red = ASD). Brackets indicate unpaired statistical comparisons (\**p* < 0.05, two-tailed permutation tests, FDR corrected). **E**, Multisensory gain as a function of age for NT (left) and ASD (right) individuals. Each datapoint represents an individual participant. **F**, Mean multisensory gain calculated with a moving window *k* of 7 years in increments of 1 year from 6–35 years for NT (blue trace) and ASD (red trace) participants. Colored error bounds indicate 95% CIs (bootstrapped). Gray shaded regions indicate significant group differences (*p* < 0.05, two-tailed permutation tests, FDR corrected). **G**, Race model violation for ASD and sex-and age-matched NT adults separated into 18–23 years (*n* = 12, left) and 24– 40 years (*n* = 11, right). **H**, Multisensory gain for the same adult groups.

While it is common practice to interpret only the positive AUC as an index of multisensory interactions (Miller, 1986; Nozawa et al., 1994; Hughes et al., 1998), equation 6 is equal to the sum of the positive and negative AUC (Colonius and Diederich, 2006; Krueger Fister et al., 2016). This is mathematically equivalent to the difference between predicted and empirical benefits, and represents the overall behavioral gain across the participant’s entire RT distribution. Qualitatively, this is equivalent to using only the positive AUC (e.g., Nidiffer et al., 2016), because the positive and negative AUCs are inversely proportional (see Fig. 4B). Moreover, the majority of younger participants in this study did not exceed statistical facilitation, rendering a statistical analysis based on the positive AUC less powerful. All race model analyses were conducted using the RaceModel open-source toolbox (https://github.com/mickcrosse/RaceModel).

### Modelling multisensory competition

Raab’s race model has been shown to provide strong predictions of empirical multisensory benefits in healthy adult participants, despite not accounting for interactions explicitly (Otto et al., 2013; Innes and Otto, 2019). We wished to determine whether multisensory benefits in children reflected interactions of a competitive or facilitative nature. Facilitation would likely follow the predictions of the race model. However, a competition could yield at least two possible scenarios: 1) the more dominant sensory modality would always win out and trigger a response or 2) the modality of the previous trial would always win out and trigger a response. Each scenario can be modelled mathematically, analogous to the race model. The first scenario was modelled with a bias towards either the auditory modality (Model 1A) or the visual modality (Model 1V) as follows:

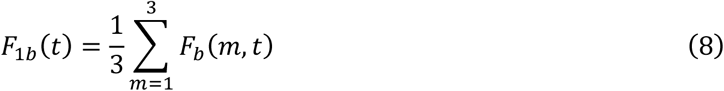

where *m* is the preceding modality (A, V, AV) and *b* is the modality that the system is biased towards (A or V). The second scenario was modelled with a bias towards the previous modality, except on AV trials where it was biased towards either the auditory modality (Model 2A) or the visual modality (Model 2V):

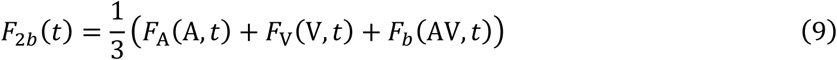

The models were used to obtain measures of predicted benefits and assessed based on how accurately they predicted empirical benefits. To examine potential developmental transitions in multisensory processing, we parametrically varied the probability of an interaction being facilitative (race model) or competitive (bias model) as follows:

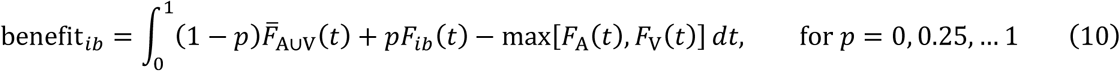

where *F*_*ib*_(*t*) is the bias model and *p* is the probability of it triggering a response. When *p* = 0, the interaction is purely facilitative (race model), and when *p* = 1, the interaction is purely competitive (bias model). In addition, Model 1 was used to examine developmental changes in sensory dominance which are likely to have a significant impact during competitive multisensory interactions.

### Modality switch effects

When testing the race model, randomly interleaving sensory modalities is necessary to minimize the opportunity for different processing strategies to be deployed under unisensory and multisensory conditions and hence satisfy the assumption of context invariance (Gondan and Minakata, 2016; Miller, 2016; Otto and Mamassian, 2017). While modality switch effects are inherent to such task conditions, their size and contribution to processes such as the RSE are rarely if ever quantified. It has been suggested that reporting the size of MSEs should become a routine procedure in RSE studies and that failure to do so would render such studies incomplete (see Otto and Mamassian, 2017). Accordingly, we measured MSEs in each participant and assessed whether or not they were likely to account for the observed RSE.

To examine MSEs, RTs were separated into those preceded by the same modality (repeat trials) and those preceded by a different modality (switch trials). Unisensory trials preceded by multisensory trials (AV→A, AV→V) were excluded from this analysis as they were considered neither switches nor repeats (repeat trials: A→A, V→V, AV→AV; switch trials: V→A, A→V, V→AV, A→AV). Separate CDFs were obtained for switch and repeat trials within each condition. Trials belonging to the two multisensory switch conditions (A→AV, V→AV) were pooled to produce one multisensory switch condition (V/A→AV). MSEs were quantified by the area between the CDFs of the switch and repeat trials:

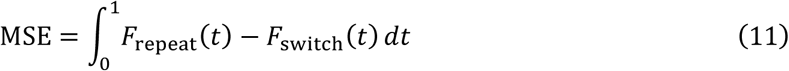

To examine the impact of switching sensory modality on the observed multisensory gain, separate tests of the race model were performed for switch and repeat trials.

### Modelling channel dependency and RT variability

It is widely considered that violation of the race model necessitates the rejection of its basic architecture in favor of the so-called coactivation model, whereby multisensory activity is pooled or integrated prior to the formation of a decision (Miller, 1982). Alternatively, sensory evidence could accumulate along separate channels that interact with one another, forming separate decisions that are then coupled by a task-relevant logical operation (Mordkoff and Yantis, 1991; Townsend and Wenger, 2004; Otto and Mamassian, 2017). Seminal work by Otto and Mamassian (2012) demonstrated that the basic race architecture can be used to explain empirical multisensory RT data by including two additional parameters to account for the additional variability or noise *η*, typically observed in empirical multisensory RTs compared to that predicted by probability summation, and the correlation *ρ* between RTs to signals on different sensory channels. Figure 8A illustrates the effect of trial history on the correlation between RTs on different channels as a function of RT quantile. Conceptually, Miller’s and Grice’s bounds assume a perfect negative and positive correlation respectively, whereas Raab’s model assumes zero correlation (i.e., independence). Otto’s context variant race model on the other hand makes no such assumptions, allowing the correlation parameter *ρ* to vary in a way that optimizes how the model predicts the empirical data.

**Figure 5.**
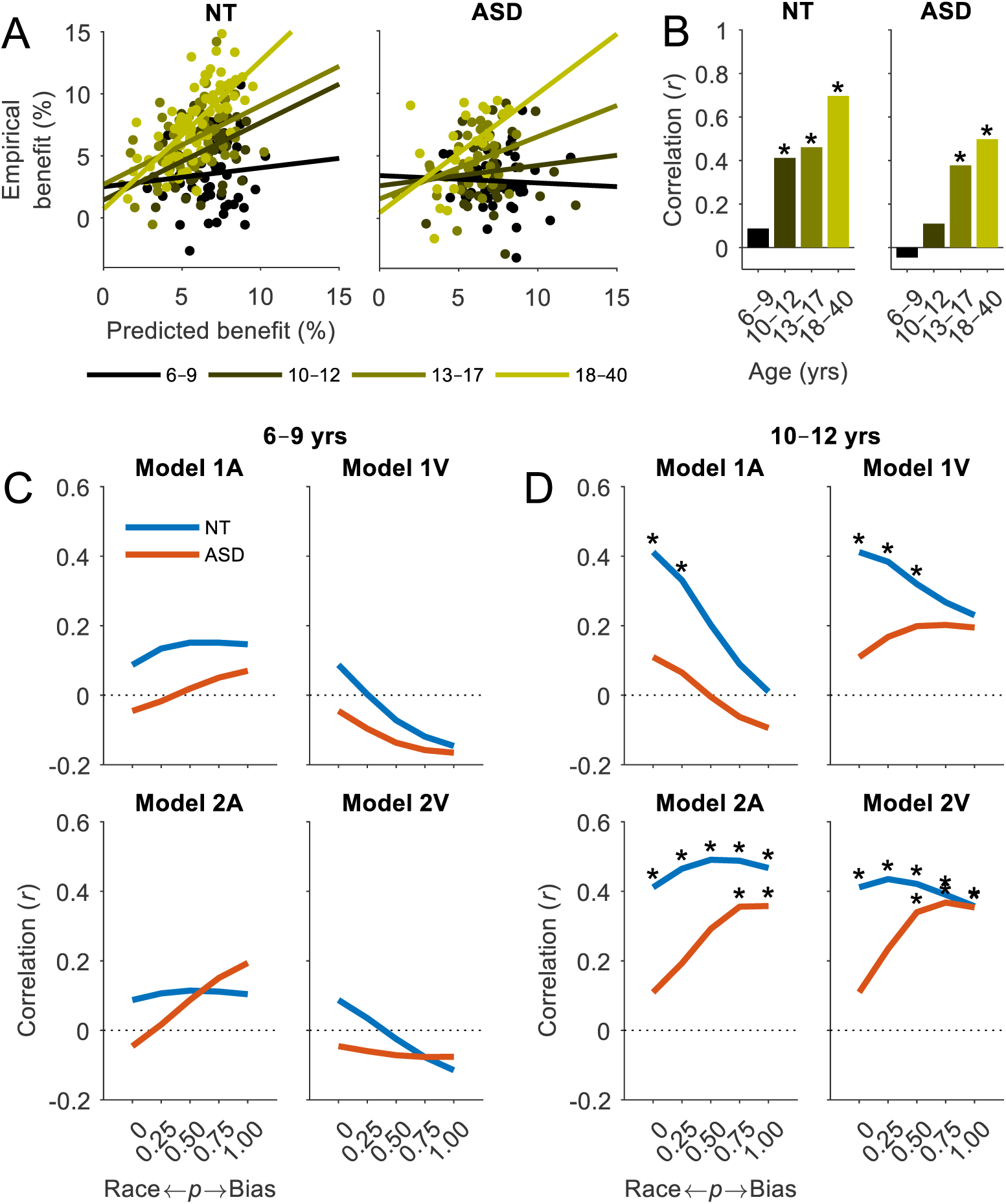
Predicting multisensory benefits. **A**, Predicted benefits versus empirical benefits for NT (left panel) and ASD (right panel) participants. Each datapoint represents an individual participant and age group is indicated by color. Solid lines represent linear fits to the data by age group. **B**, Pearson correlation coefficient (*r*) of the regression fits in panel A. Asterisks indicate significant correlations (*p* < 0.05, two-tailed permutation tests). **C**, **D**, Hypothetical models of multisensory competition were tested. Model 1A was biased towards the auditory modality and Model 1V towards the visual modality (Eq. 8). Model 2A was biased towards the preceding modality and the A modality when preceded by an AV trial, and Model 2V was biased towards the preceding modality and the V modality when preceded by an AV trial (Eq. 9). The probability *p* of an interaction being facilitative (race model) or competitive (bias model) was parametrically varied between 0 and 1 in increments of 0.25 (Eq. 10). The ability the models to predict empirical benefits was assessed within each age group based on the Pearson correlation coefficient. Data presented are the two younger age groups. See supplementary material for the two older age groups.

**Figure 6.**
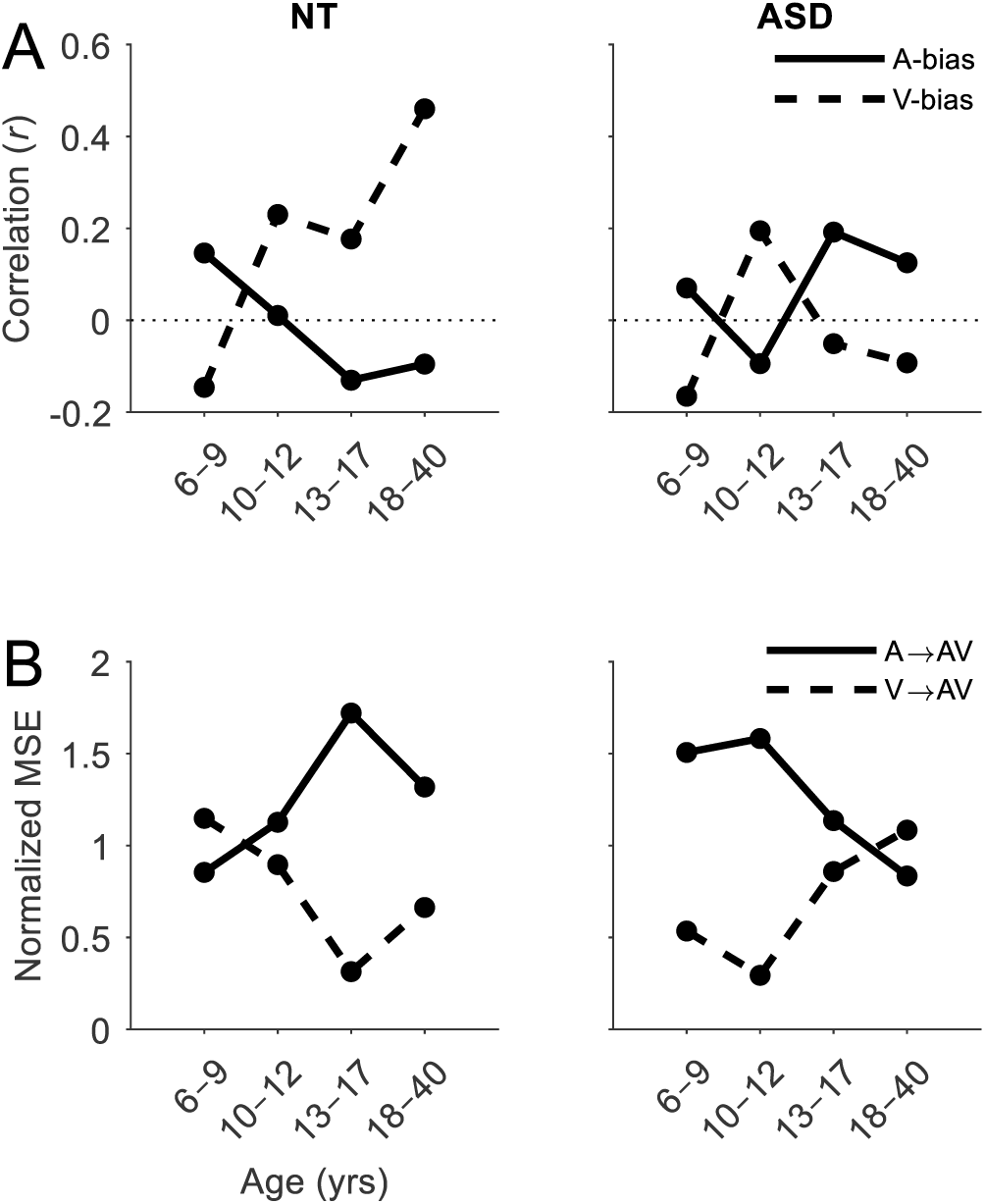
Developmental changes in sensory dominance. **A**, Sensory dominance was examined by measuring the performance of Model 1A (A-bias, solid trace) and Model 1V (V-bias, dotted trace) with the probability of a sensory-specific bias *p* set to 1 (Eq. 10). The ability of each model to predict empirical benefits was assessed within each age group based on the Pearson correlation coefficient. **B**, Modality switch effects for AV trials separated by trials preceded by A-stimuli (solid trace) and V-stimuli (dotted trace). MSEs were quantified by the area between the CDFs of the switch and repeat trials and normalized by the grouped V/A→AV MSEs.

**Figure 7.**
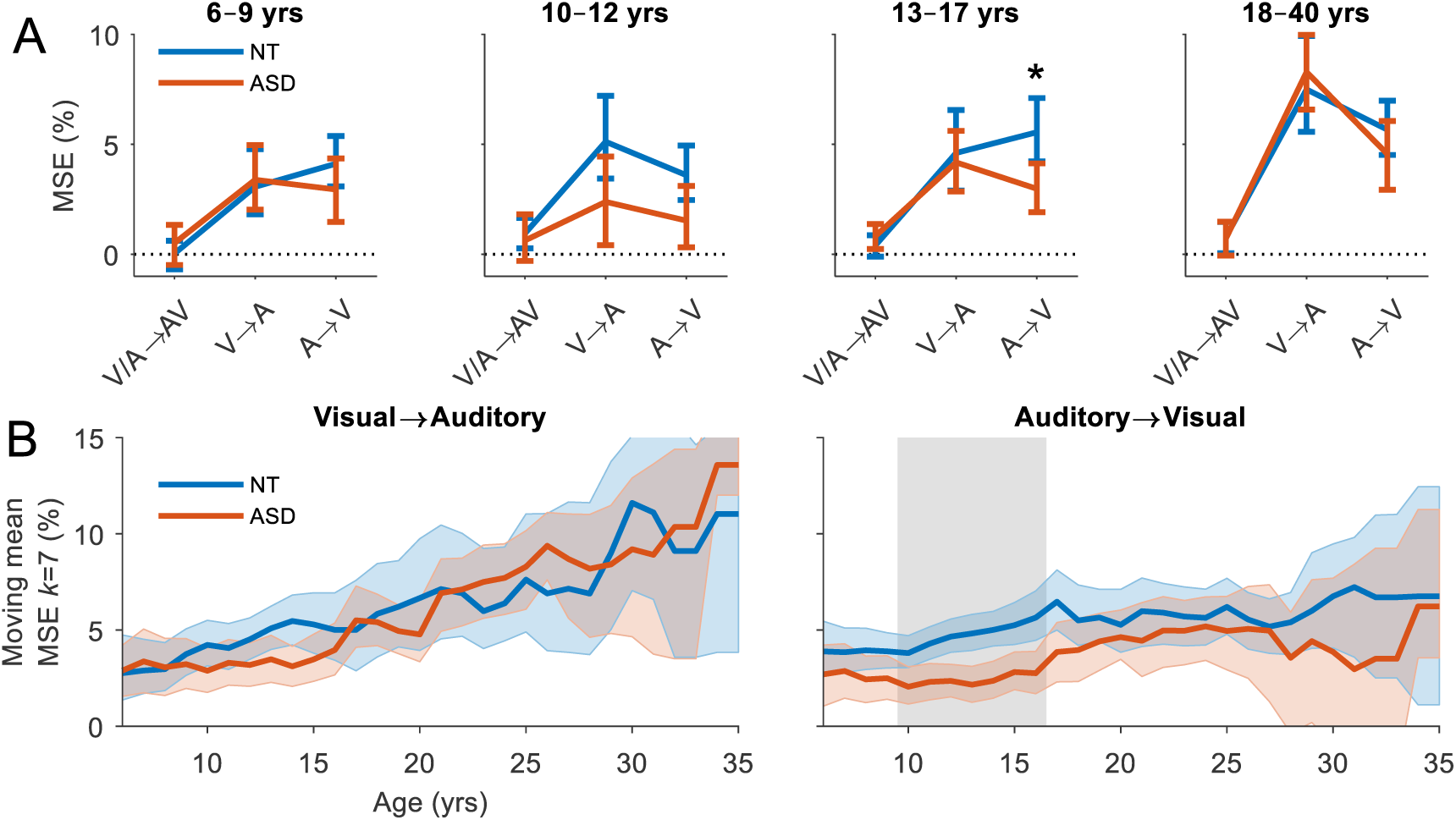
Modality switch effects. **A**, Mean MSE for each condition by age group. MSEs were quantified by the area between the CDFs of the switch and repeat trials (Eq. 8). Error bars indicate 95% CIs (bootstrapped). Asterisks indicate significant group differences (*p* < 0.05, two-tailed permutation tests, *t*_max_ corrected). **B**, Mean MSE for visual to auditory (left panel) and auditory to visual (right panel) switches calculated with a moving window *k* of 7 years in increments of 1 year from 6–35 years for NT (blue trace) and ASD (red trace) participants. Colored error bounds indicate 95% CIs (bootstrapped). Gray shaded regions indicate significant group differences (*p* < 0.05, two-tailed permutation tests, FDR corrected).

**Figure 8.**
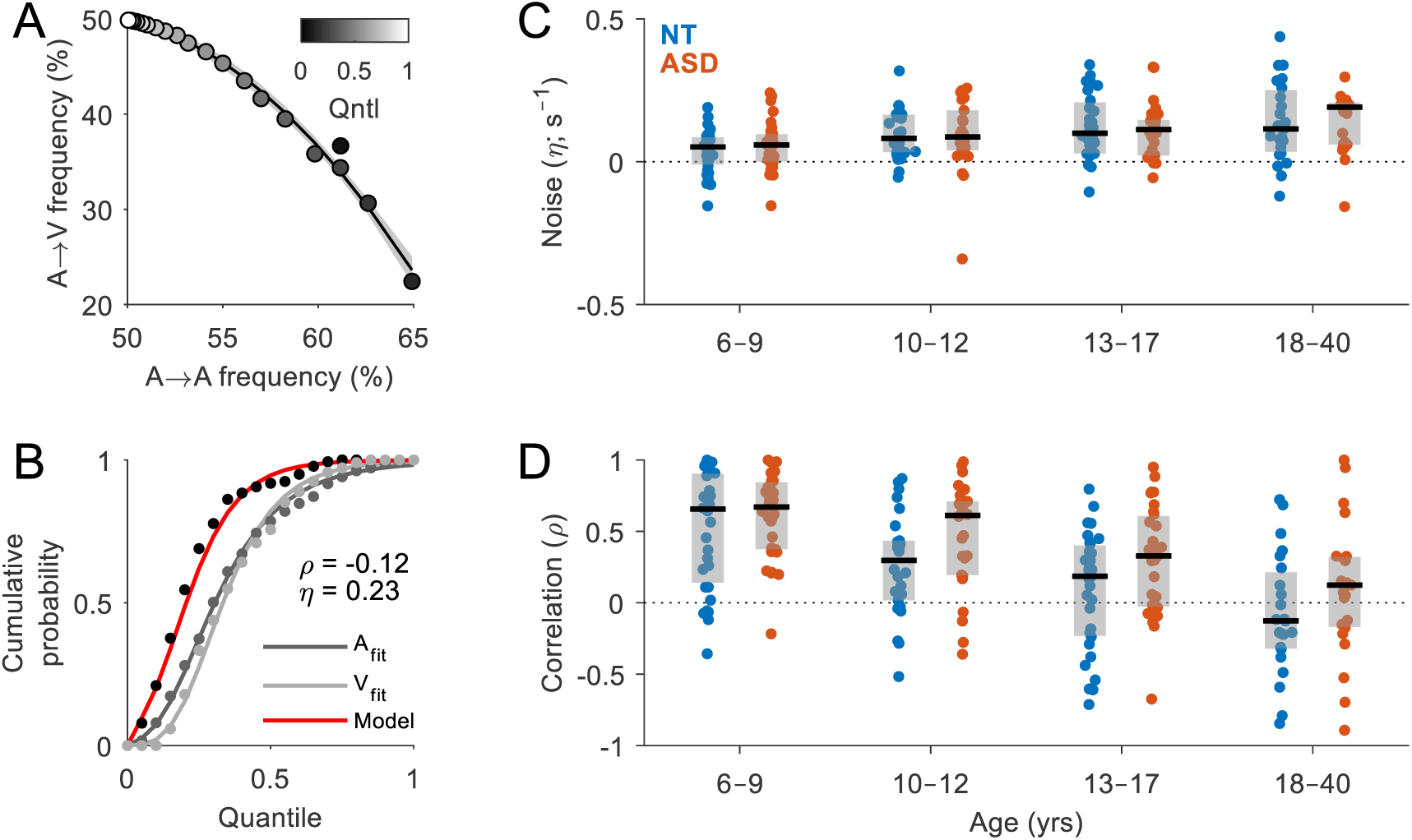
Modelling channel dependency and RT variability. **A**, Frequency of visual and auditory trials preceded by auditory trials in each quantile (i.e., switch versus repeat trials). Quantiles are indicated by a grayscale, graduating from black (fastest quantile) to white (slowest quantile). Example data averaged over all NT adult participants. **B**, CDFs were fit to the unisensory RT data and used to predict empirical multisensory RT data via Otto’s context variant race model (Otto and Mamassian, 2012). Free parameters *ρ* and *η* account for the correlation between RTs on different channels and increased RT variability or noise, respectively. Data from an example NT adult participant. **C**, **D**, Best-fitting model parameters *ρ* and *η* by diagnosis and age group. Boxplots indicate the median value (black line) and interquartile range (grey box). Each datapoint represents an individual participant (blue = NT, red = ASD).

Applying this modelling approach, we examined the values of *ρ* and *η* that optimized the model fit for each participant in order to gain additional insight into the cognitive processes underlying group differences in multisensory processing and modality switching. Using the RSE-box (v1.0) toolbox (https://github.com/tomotto/RSE-box; Otto, 2018), Gaussian functions were fit to the reciprocal of the unisensory RT distributions via the LATER model approach (Noorani and Carpenter, 2016), which assumes that the reciprocals of the RT distributions are normally distributed with mean *μ* and SD *σ* (see Fig. 8B). These parameters were then used to generate the probability density function (PDF) of the maximum distribution *f*_A∪V_(x) = *f*_A_(-x) + *f*_V_(-x), where

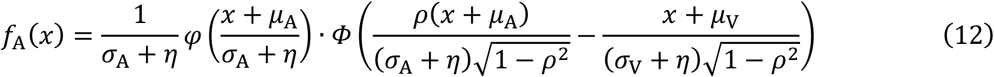

where *φ* and *Φ* are the PDF and CDF of the standard normal distribution, respectively. Calculation of *f*_V_(x) was obtained analogously to equation 9. A more detailed description can be found in Otto and Mamassian (2012), supplementary information.

### Statistical analyses

As an initial inquiry, a linear mixed-effects model was used to determine which parameters influenced RTs. The model was fit using the maximum likelihood criterion. Single-trial RTs were the continuous numeric dependent variable. Diagnosis was a contrast-coded fixed factor (NT, ASD), age was a continuous numeric fixed factor (6–40 years), and condition was a multi-level nominal fixed factor (AV, A, V). Subjects were included as a random factor, along with by-subject slope adjustments for condition (Barr et al., 2013). ISI was included as another random factor, as well as preceding modality with slope adjustments for condition. Subsequent analyses employing standard linear models coded fixed effects as above. A one-way analysis of covariance (ANCOVA) was used to assess the correspondence between empirical and predicted benefits, treating age group as a partialled out categorical variable (Bland and Altman, 1995).

A mediation analysis (Baron and Kenny, 1986) was used to establish whether the relationship between participants’ age and multisensory gain was mediated by a direct effect of age on MSE. Age was chosen as the causal variable in the model because of its known effect on race model violation (Brandwein et al., 2011). For this analysis, MSEs were averaged across the two unisensory conditions (V→A, A→V), as we hypothesized that it was a slowing of unisensory RTs that was the cause of the observed RSE. Using the M3 Toolbox (https://github.com/canlab/MediationToolbox), we constructed a three-variable mediation model with age as the causal variable, gain as the outcome variable and MSE as the mediating variable (Fig. 9C). For MSE to be considered a mediator, the following criteria must be met based on three separate regressions: 1) the causal variable must affect the outcome, 2) the causal variable must affect the mediator, and 3) the mediator must affect the outcome but the causal variable must either no longer affect the outcome (full mediation) or at least weaken the effect (partial mediation). Significance and SE of the associated path coefficients were bootstrapped (10,000 samples) and adjusted using the bias-corrected and accelerated percentile method (Wager et al., 2008).

**Figure 9.**
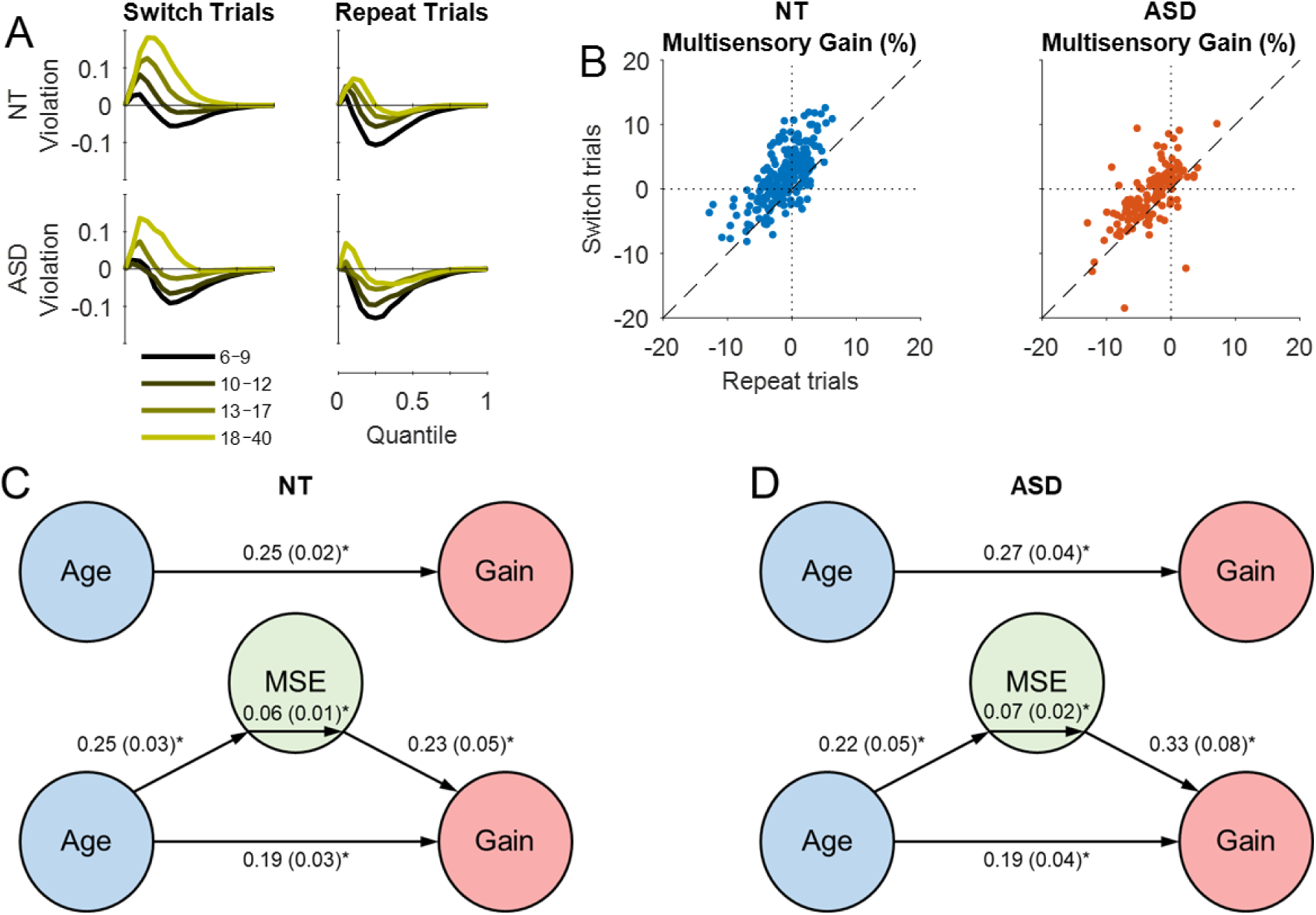
Linking modality switch effects and redundant signals effects. **A**, Race model violation by diagnosis and age for switch trials (left panel) and repeat trials (right panel). **B**, Multisensory gain on switch trials versus repeat trials for NT (left panel) and ASD (right panel) individuals. Each datapoint represents an individual participant. **C**, **D**, Mediation model that tested whether modality switch effects (MSEs) mediated the effect of age on multisensory gain. Paths between nodes are labeled with regression coefficients, with SE in parentheses (\**p* < 0.001, bootstrapped). In both groups, age predicted gain (top path), and predicted MSE controlling for gain (lower left path). The middle coefficients indicate formal mediation effects but the significant direct paths between age and gain controlling for MSE (bottom path) suggest only partial mediation, i.e., MSE did not explain all of the shared variance between age and gain.

All *post hoc* statistical comparisons were conducted using nonparametric permutation tests (10,000 permutations) based on the *t*-statistic and adjusted to control for family-wise error rate using the *t*_*max*_ correction method (Westfall and Young, 1993; Blair et al., 1994). This method has been shown to control for Type 1 error at a desired level when performing tests of the race model at multiple quantiles and the power of the test is reasonable even for small samples (Gondan, 2010). Equivalence of variance was established prior to all unpaired tests using a permuted *F*-test and the appropriate *t*-statistic was then applied based on the outcome. Effect sizes were calculated using Cohen’s *d* and were bias-corrected according to sample size (Hedges and Olkin, 1985). All confidence intervals (CIs) were bootstrapped (10,000 samples) at the 95% confidence level and adjusted using the bias-corrected and accelerated percentile method (Davison and Hinkley, 1997). Correlation analyses were conducted using permuted Pearson correlation or Spearman rank coefficients (Bishara and Hittner, 2012). All *post hoc* statistical tests and effect size calculations were conducted using the PERMUTOOLS open-source toolbox (https://github.com/mickcrosse/PERMUTOOLS).

## Results

### Reaction times and multisensory benefits

A linear mixed-effects analysis was used to examine the effect of diagnosis, age and stimulus condition on response times (*R*^2^_adj_ = 0.495). Subjects, ISI and preceding modality were included as random factors, along with slope adjustments for condition (see Methods for details). Participants with ASD responded more slowly to stimuli than their NT peers (*β* = 47.6, *SE* = 12.3, *p* = 0.0001; Fig. 2A). There was an effect of maturation, with older participants responding faster than younger participants (*β* = −9.1, *SE* = 0.84, *p* = 2×10^-27^). Responses to multisensory stimuli were faster than those to both audio (*β* = 55.2, *SE* = 9.96, *p* = 3×10^-8^) and visual (*β* = 67.1, *SE* = 6.7, *p* = 6×10^-24^) stimuli, indicating the presence of an RSE. There was an interaction between age and RSE (RSE_A_: *β* = −0.6, *SE* = 0.22, *p* = 0.006; RSE_V_: *β* = −0.48, *SE* = 0.18, *p* = 0.008).

To examine the RSE in detail, a general linear model was constructed to quantify the effects of diagnosis and age on predicted (*R*^2^_adj_ = 0.086) and empirical benefits (*R*^2^_adj_ = 0.255). Predicted benefits decreased as a function of age (*β* = −0.07, *SE* = 0.01, *p* = 1×10^-7^) and were not significantly different in NT and ASD individuals (*β* = 0.3, *SE* = 0.2, *p* = 0.12). Conversely, empirical benefits increased with age (*β* = 0.18, *SE* = 0.02, *p* = 2×10^-17^) and were smaller in ASD individuals (*β* = −1.5, *SE* = 0.3, *p* = 7×10^-7^). This suggests that the race model over-predicts empirical benefits for younger individuals and under-predicts them for older individuals (see Fig. 2C). Moreover, the race model does not predict the group differences in empirical multisensory benefits, suggesting an integrative deficit.

### Testing the race model

To determine whether the RSE exceeded statistical facilitation, we compared the multisensory CDFs to the race model at each of the first 7 quantiles (maximum number of quantiles violated by any group). Violation of the race model was assessed using right-tailed permutation tests with *t*_max_ correction (Gondan, 2010). NT participants showed evidence of violation at one or more quantiles in every age group, the number of quantiles increasing as a function of age (*p* < 0.05, shaded area, Fig. 3A). The percentage of participants that exceeded statistical facilitation at each quantile is illustrated in Figure S1. Individuals with ASD showed no evidence of violation between the ages of 6–12 years (Fig. 3B). However, evidence of violation emerges in adolescence (first quantile) and becomes more evident in adulthood (first 2 quantiles; see Table 2 for the statistics of each race model test). Note, these results were replicated qualitatively using the more conservative Miller’s bound, albeit at less quantiles (see Table S1).

**Table 2.** Test statistics comparing CDFs of multisensory RTs with the race model. Values shown indicate effect sizes (Cohen’s *d* corrected for sample size) and 95% CIs (bootstrapped) in brackets. Asterisks indicate significant race model violation (*p* < 0.05, right-tailed permutation tests, *t*_max_ corrected).

| Q | NT |  |  |  | ASD |  |  |  |
| --- | --- | --- | --- | --- | --- | --- | --- | --- |
|  | 6–9 yrs | 10–12 yrs | 13–17 yrs | 18–40 yrs | 6–9 yrs | 10–12 yrs | 13–17 yrs | 18–40 yrs |
| 1 | 0.18[0.1,0.3]* | 0.44[0.3,0.7]* | 0.54[0.4,0.9]* | 0.83[0.6,1.1]* | 0.12[-0.0,0.3] | 0.07[-0.1,0.3] | 0.30[0.1,0.6]* | 0.62[0.4,1.1]* |
| 2 | -0.06[-0.2,0.1] | 0.21[0.1,0.4]* | 0.41[0.3,0.6]* | 0.73[0.6,0.9]* | -0.14[-0.2,-0.0] | -0.06[-0.2,0.1] | 0.09[-0.1,0.3] | 0.38[0.2,0.6]* |
| 3 | -0.21[-0.3,-0.1] | 0.03[-0.1,0.2] | 0.29[0.2,0.4]* | 0.59[0.4,0.8]* | -0.29[-0.4,-0.2] | -0.19[-0.3,-0.1] | -0.08[-0.3,0.1] | 0.19[0.0,0.4] |
| 4 | -0.35[-0.5,-0.2] | -0.11[-0.2,-0.0] | 0.17[0.1,0.3]* | 0.44[0.3,0.6]* | -0.48[-0.7,-0.3] | -0.34[-0.6,-0.2] | -0.29[-0.6,-0.1] | 0.07[-0.1,0.3] |
| 5 | -0.53[-0.8,-0.4] | -0.27[-0.4,-0.2] | 0.03[-0.1,0.2] | 0.31[0.2,0.4]* | -0.66[-1.1,-0.5] | -0.50[-0.8,-0.3] | -0.54[-0.9,-0.3] | 0.01[-0.2,0.2] |
| 6 | -0.80[-1.1,-0.6] | -0.43[-0.7,-0.3] | -0.09[-0.2,0.0] | 0.21[0.1,0.3]* | -0.89[-1.4,-0.6] | -0.70[-1.2,-0.5] | -0.75[-1.3,-0.5] | -0.07[-0.3,0.1] |
| 7 | -1.00[-1.4,-0.8] | -0.55[-0.8,-0.4] | -0.20[-0.4,-0.1] | 0.12[-0.0,0.2] | -1.12[-1.7,-0.8] | -0.75[-1.3,-0.6] | -0.97[-1.5,-0.6] | -0.17[-0.4,-0.0] |

To compare race model violation between NT and ASD individuals of different ages, we computed the root-mean-square error (RMSE) and correlation coefficient between each participant’s violation function and that of every other participant. Because the violation functions are typically non-normal, we applied a rank-based inverse normal (RIN) transformation (Bliss, 1967), prior to assessing the Pearson correlation. Participants were split into 8 age groups separated by 3 years between the ages of 6–30 years (there were too few participants above 30 years of age). Matrices containing RMSE and correlation values were obtained by averaging over the values within each age group (Fig. 3C). The red line in Figure 3C indicates the age groups that are most similar, and its divergence above the dotted midline suggests that multisensory behavior in ASD participants corresponded more closely to that of younger NT participants, i.e., a developmental delay. Convergence of the red and dotted lines suggests that this delay may recover in adulthood, in line with our original hypothesis. This is further examined in the following section.

### Delayed multisensory development in autism

We constructed a linear model to evaluate the effects of diagnosis and age on multisensory gain (*R*^2^_adj_ = 0.388). Multisensory gain, as indexed by the AUC (Eq. 6, Fig. 4A), increased as a function of age (*β* = 0.25, *SE* = 0.02, *p* = 2×10^-21^) but was significantly reduced in participants with ASD compared to NT individuals (*β* = −1.98, *SE* = 0.68, *p* = 0.004). The absence of an interaction suggests that this maturation effect was present in both groups (*β* = 0.01, *SE* = 0.04, *p* = 0.77). *Post hoc* comparisons were conducted within each of the four age groups. For this analysis, NT participants were sex-and age-matched to each of the ASD participants and compared at every quantile using two-tailed (unpaired) permutation tests. Group differences were observed in the adolescent group at quantiles 4 and 5 (*p* < 0.05, shaded area, Fig. 4C). To compare the overall multisensory gain, we conducted permutation tests on the AUC (Fig. 4D), revealing differences in participants aged 10–12 years (*t*_(50)_ = 2.22, *p* = 0.031, *d* = 0.61, 95CI [0.1, 1.15]) and 13–17 years (*t*_(60)_ = 2.57, *p* = 0.014, *d* = 0.65, 95CI [0.18, 1.15]), but not 6–9 years (*t*_(60)_ = 0.88, *p* = 0.39, *d* = 0.21, 95CI [-0.28, 0.72]) or 18–40 years (*t*_(44)_ = 1.81, *p* = 0.077, *d* = 0.52, 95CI [-0.03, 1.19]). The moderate effect size in the adult group suggests that individuals with ASD might not have “caught up” entirely by 18 years of age.

The effect of maturation can be seen more clearly by charting multisensory gain as a function of age (Fig. 4E). Age was highly predictive of multisensory gain between 6–17 years (NT: *R*^2^ = 0.34, *p* = 0; ASD: *R*^2^ = 0.21, *p* = 0) but not between 18–40 years (NT: *R*^2^ = 0.005, *p* = 0.56; ASD: *R*^2^ = 0.052, *p* = 0.296), suggesting that maturation of this process ceases in adulthood. To characterize this developmental trajectory more precisely, we calculated the mean multisensory gain with a moving window *k* of 7 years in increments of 1 year (Fig. 4F). Controls were sex-and age-matched to ASD individuals within each 7-year window and compared using two-tailed permutation tests (FDR corrected). In NT participants, multisensory gain increased steadily between 6–18 years of age. In individuals with ASD, the rate of increase was more gradual and was significantly lower than that of their NT peers between the ages of 11–21 years (*p* < 0.05, shaded area, Fig. 4F). However, by the mid-twenties, multisensory gain was commensurate with that of NT individuals suggesting that this deficit recovers in adulthood, confirming our original hypothesis. Given that maturation appears to continue well into adulthood, a *post hoc* analysis was conducted whereby the adult group was subdivided into participants aged 18–23 years (*n* = 12) and 24–40 years (*n* = 11) to examine multisensory gain before and after this “catch up” point. As expected, there were significant group differences in adults aged 18–23 years (*t*_(20)_ = 2.24, *p* = 0.039; *d* = 0.92, 95CI [0.18, 1.98]; Fig. 4H, left) but not in adults aged 24–40 years (*t*_(22)_ = 0.36, *p* = 0.72; *d* = 0.14, 95CI [-0.64, 1.0]; Fig. 4H, right). Group average violation functions were almost identical at every quantile between NT and ASD adults aged 24–40 years (Fig. 4G, right), suggesting both qualitative and quantitative recovery.

### Predicting multisensory benefits

Raab’s race model has been shown to provide overall a strong prediction of the RSE in healthy adults (Otto et al., 2013; Innes and Otto, 2019). To assess whether the race model could predict multisensory benefits in children and individuals with ASD, one-way ANCOVAs were used to measure the correlation between predicted and empirical benefits in each age group. Predicted benefits were correlated with empirical benefits in both NT individuals (*F*_(1,217)_ = 62.86, *p* = 1×10^-13^, *R*^2^ = 0.23) and individuals with ASD (*F*_(1,125)_ = 5.25, *p* = 0.024 *R*^2^ = 0.04) but an interaction suggested that this relationship was age-dependent (NT: *F*_(3,217)_= 5.4, *p* = 0.0013, *R*^2^ = 0.07; ASD: *F*_(3,125)_ = 2.58, *p* = 0.057, *R*^2^ = 0.06). Figure 5A, B shows that the ability of the race model to predict empirical benefits increases dramatically as a function of age. While the race model predicted a significant proportion of the variance in the adult groups (NT: *R*^2^ = 0.49, *p* = 0; ASD: *R*^2^= 0.25, *p* = 0.017), it accounted for almost none of the variance in the youngest groups (NT: *R*^2^ = 0.007, *p* = 0.55; ASD: *R*^2^ = 0.002, *p* = 0.78).

To determine whether multisensory benefits in children reflected interactions of a competitive nature, we tested the predictions of models that were biased towards either a specific modality (Model 1A, V) or the previous modality (Model 2A, V). The probability *p* of an interaction being facilitative (race model) or competitive (bias model) was parametrically varied between 0 and 1 in increments of 0.25. In children with ASD aged 6–9 years, Model 2A was most accurate at predicting empirical benefits, suggesting that their responses were triggered by the previous modality, with a bias towards the auditory cue (Fig. 5C). In their NT peers, none of the bias models exceeded the performance of the race model considerably, but there was evidence for an auditory bias once again. In ASD participants aged 10–12 years, Model 2V dramatically exceeded race model performance, suggesting that RTs were largely determined by the previous modality, but this time, with a bias towards the visual modality (Fig. 5D). In their NT peers, there was no major improvement beyond the race model, but there was evidence of a visual bias as well. In teenagers and adults, none of the bias models outperformed the race model, suggesting that individuals with ASD transition from competition to facilitation in adolescence (Fig. S2).

### Developmental changes in sensory dominance

To examine developmental patterns in sensory dominance, we tested Model 1A and Model 1V with the probability of a sensory-specific bias set to 1 (Eq. 10). Evaluating model performance as before, we noticed an auditory dominance in both groups at 6–9 years of age that shifted to a visual dominance by 10–12 years of age (Fig. 6A). In the NT group, this visual dominance appears to continue into adulthood in accordance with the well-known Colavita visual dominance effect (Colavita, 1974). However, in the ASD group, this sensory weighting appears to shift once again in adolescence, leading to an auditory dominance in adulthood.

If such sensory dominances exist, one would expect to see greater modality switch effects (MSEs) on AV trials preceded by the less dominant modality than those preceded by the dominant modality. To test this hypothesis, we examined MSEs on AV trials, separating trials preceded by A and V stimuli (A→AV and V→AV, respectively). MSEs were quantified by deriving separate CDFs for switch and repeat trials and computing the area between them (Eq. 11). Additionally, MSEs in each condition were normalized by that of the grouped condition (V/A→AV) to allow for meaningful comparison across age groups (this did not change the results qualitatively). Based on our modelling analysis, we expected to see greater MSEs on V→AV trials for TD children (6–9 years) and on A→AV trials for older TD children and adults (10–40 years). We expected something similar for ASD individuals with another shift in adolescence. The data in Figure 6B suggest that, as predicted, MSEs were greater on V→AV trials for TD children (6–9 years) and on A→AV trials for older TD children and adults (10–40 years). For ASD individuals, the data suggest the reverse, with greater MSEs on A→AV trials in children and teenagers (6–17 years) and on V→AV trials for adults (18–40 years).

### Modality switch effects

We modelled the effects of diagnosis, age and condition on MSEs using a linear model (*R*^2^_adj_ = 0.303). MSEs increased with age (*β* = 0.17, *SE* = 0.02, *p* = 7×10^-24^) and were reduced in individuals with ASD compared to NT individuals (*β* = −1.18, *SE* = 0.24, *p* = 7×10^-7^). Compared to multisensory trials, MSEs were larger on both auditory trials (*β* = 4.67, *SE* = 0.03, *p* = 6×10^-57^) and visual trials (*β* = 3.66, *SE* = 0.03, *p* = 3×10^-37^). Follow-up permutation tests revealed that MSEs were only reduced in the adolescent ASD group, and only when switching from auditory to visual stimuli (*t*_(60)_ = 2.76, *p* = 0.021, *d* = 0.69, 95CI [0.22, 1.19]; Fig. 7A). A more detailed examination using a moving mean estimate of MSE showed that group differences emerged between the ages of 10–16 years (*p* < 0.05, shaded area, Fig. 7B, right). The maturational course of visual to auditory MSEs appears to continue later into development than that of auditory to visual switches in both groups (Fig. 7B, left).

Contrary to our results, a study by Williams et al. (2013) found that individuals with ASD between the ages of 8–15 years exhibited a greater cost to switching from auditory to visual stimuli than their age-matched NT peers. To make a more direct comparison with their study, we performed a two-tailed permutation test on a group of sex-and age-matched participants between the ages of 8–15 years (*n* = 72) and used a similar measure of MSE based on mean RT values. This approach yielded the same outcome as before, with ASD individuals exhibiting smaller MSEs (NT: 30.5 ± 27.4 ms, ASD: 19.7 ± 38.7 ms; *t*_(142)_ = 1.93, *p* = 0.049, *d* = 0.32, 95CI [-0.004, 0.66]), confirming the discrepancy was not the result of how MSE was quantified. The only remaining difference between our two studies was that Williams et al. (2013) used longer ISIs (3–5 s versus 1–3 s). Thus, we repeated the test focusing on RTs with preceding ISIs between 2.5–3 s. Limiting the analysis to longer ISIs caused a significant drop in MSE for NT individuals (16.2 ± 42.8 ms) but not so much for individuals with ASD (16.8 ± 51.5 ms). Moreover, this modification revealed no group differences (*t*_(142)_ = −0.07, *p* = 0.954, *d* = −0.01, 95CI [-0.34, 0.31]), suggesting invocation of disparate mechanisms underlying MSEs at shorter versus longer ISIs.

### Increased channel dependency in autism

To gain a better understanding of what aspects of multisensory processing led to differences in behavior, we adopted a computational framework based on the race model (Otto and Mamassian, 2012). The inclusion of 2 additional free parameters in the race model allowed us to quantify the additional variability or noise *η* in empirical multisensory RTs, as well as the correlation *ρ* between RTs on different sensory channels, giving us insight into how attention is divided between them (see Methods for details). We hypothesized that the increase in RT variability would be larger for individuals with higher multisensory gain, and that channel dependency would be lower or more negatively correlated for individuals with greater MSEs. The best-fitting estimates of the noise parameter *η* increased with age (*β* = 0.0045, *SE* = 0.0008, *p* = 4×10^-8^) but was not statistically different between NT and ASD participants (*β* = −0.016, *SE* = 0.012, *p* = 0.17; *R*^2^_adj_ = 0.0899; Fig. 8C). The best-fitting estimates of the correlation parameter *ρ* decreased with age (*β* = −0.035, *SE* = 0.003, *p* = 1×10^-29^) and were lower (and sometimes more negative) for NT individuals (*β* = −0.25, *SE* = 0.04, *p* = 2×10^-9^; *R*^2^_adj_ = 0.38; Fig. 8D). *Post hoc* permutation tests revealed moderate group differences in participants aged 10–12 years (*t*_(50)_ = 1.97, *p* = 0.05, *d* = 0.54, 95CI [0.01, 1.18]) and 13–17 years (*t*_(60)_ = 2.15, *p* = 0.036, *d* = 0.54, 95CI [0.06, 1.06]). This greater (more positive) channel dependency in ASD suggests a more even spread of attention across sensory systems.

### Linking modality switch effects and redundant signals effects

To examine the relationship between MSEs and multisensory gain, we performed a series of partial correlations across participants, controlling for age (Table 3). As one might predict, there was a strong positive correlation between the average multisensory gain on switch trials and the average MSE on unisensory trials (but not on multisensory trials). However, there was no significant correlation between multisensory gain on repeat trials and MSEs on unisensory trials, whereas there was a strong positive correlation with MSEs on multisensory trials. This pattern, which was identical in both groups (see Fig. S3), confirms that MSEs on unisensory trials are more likely to contribute to multisensory gain. Figure 9A, B illustrates the impact of switching sensory modality on race model violation and multisensory gain, respectively. 87% of NT individuals exhibited a larger multisensory gain on switch trials than on repeat trials (*t*_(224)_ = 15.62, *p* = 0, *d* = 0.84, 95CI [0.73, 0.96]), with 82% of individuals with ASD showing the same (*t*_(132)_ = 6.74, *p* = 0, *d* = 0.51, 95CI [0.35, 0.68]). Nevertheless, when we submitted RTs from the repeat trials to a race model test, every group violated the race model as before except the adolescent ASD group (Table S2), even when using a more conservative test based on Miller’s bound (Table S3).

**Table 3.** Partial correlations between multisensory gain and MSEs, controlling for age. Multisensory gain was computed separately for switch trials (left columns) and repeat trials (right columns). Values indicate coefficients of determination (*R*^2^) and significance of the correlation (*p*).

|  | Gain on Switch Trials |  |  | Gain on Repeat Trials |  |  |
| --- | --- | --- | --- | --- | --- | --- |
|  | V/A→AV | V→A | A→V | V/A→AV | V→A | A→V |
| NT | $R^2 = 0.001$<br>$p = 0.7$ | $R^2 = 0.3$<br>$p = 3 \times 10^{-19}$ | $R^2 = 0.18$<br>$p = 3 \times 10^{-11}$ | $R^2 = 0.25$<br>$p = 1 \times 10^{-15}$ | $R^2 = 2 \times 10^{-6}$<br>$p = 0.98$ | $R^2 = 0.003$<br>$p = 0.43$ |
| ASD | $R^2 = 0.035$<br>$p = 0.03$ | $R^2 = 0.29$<br>$p = 2 \times 10^{-11}$ | $R^2 = 0.21$<br>$p = 4 \times 10^{-8}$ | $R^2 = 0.31$<br>$p = 3 \times 10^{-12}$ | $R^2 = 0.026$<br>$p = 0.06$ | $R^2 = 0.001$<br>$p = 0.75$ |

Having established the relationship between MSEs and multisensory gain, we wished to determine whether the contribution of the former was a full or partial. To do this, we submitted the data to a mediation analysis (Wager et al., 2008). Specifically, we tested whether MSEs mediated the relationship between participant age and multisensory gain (Fig. 9C, D). First, we established that age was a reliable predictor of both MSE (NT: *β* = 0.25, *SE* = 0.03, *p* = 0.0002; ASD: *β* = 0.22, *SE* = 0.05, *p* = 0.001) and multisensory gain (NT: *β* = 0.25, *SE* = 0.02, *p* = 0.001; ASD: *β* = 0.27, *SE* = 0.04, *p* = 0.0002), meeting the first two criteria for mediation (see Methods for details). MSE affected gain, controlling for age (NT: *β* = 0.23, *SE* = 0.05, *p* = 0.0002; ASD: *β* = 0.33 *SE* = 0.08, *p* = 0.0001) and the mediation effect was significant for both groups (NT: *β* = 0.06, *SE* = 0.01, *p* = 0.0002; ASD: *β* = 0.07 *SE* = 0.02, *p* = 0.0001). However, there was still a significant direct path between age and gain when controlling for MSE (NT: *β* = 0.19, *SE* = 0.03, *p* = 0.0002; ASD: *β* = 0.19 *SE* = 0.04, *p* = 0.0004), indicating that MSE only partially mediated the observed relationship between age and multisensory gain.

## Discussion

Our data suggest that the amelioration of multisensory deficits in ASD generalizes to the case of nonsocial AV stimuli, but that the developmental trajectory of this recovery is protracted compared to that observed in AV speech studies (e.g., Taylor et al., 2010; Foxe et al., 2015). We hypothesized that this delay may be due to lack of environmental exposure to such ecologically-rare stimuli (Beker et al., 2017; Cuppini et al., 2017), and/or engagement of neural processes with longer developmental trajectories. Indeed, multisensory gain in NT individuals has been shown to reach full maturity much later for simple AV stimuli, such as those used here (Brandwein et al., 2011), compared to AV speech stimuli (Ross et al., 2011). This undoubtedly affects the average age at which we observe developmental recovery in ASD, suggesting that it is important to consider the maturational course in typically-developing individuals within different contexts when examining developmental recovery in clinical populations.

The disparity in multisensory development for speech and non-speech stimuli likely reflects the fact that multisensory processing occurs across distributed networks and that different stimuli and tasks tap into unique processes with varying maturational courses (Chandrasekaran, 2017). The task employed in the current study required the speeded detection of simple AV stimuli, without discrimination, identification or any higher-order cognitive processing. Integration of such simple AV stimuli likely consists of early cross-sensory activation of visual and auditory cortical regions, enhancing detection of the incoming visual and auditory inputs, respectively (Molholm et al., 2002; Mercier et al., 2013; Mercier et al., 2015). In contrast, identification of AV speech engages an extensive network of hierarchically-organized brain areas (Hickok and Poeppel, 2007; Peelle, 2019), mapping spectrotemporal representations to phonetic representations, and from there to lexical-semantic representations. Moreover, integration of auditory and visual speech cues may act through multiple integrative mechanisms, including early visual activation of auditory cortex, increasing perceptual sensitivity (Mégevand et al., 2018), and later integration of visual speech content (i.e., place and/or manner of articulation), reducing the density of phonemic and lexical neighborhoods (Tye-Murray et al., 2007; Peelle and Sommers, 2015). Clearly, task demands and stimuli play a major role in the patterns of multisensory deficits and recovery functions that are observed for any given experimental paradigm.

Alternatively, differences in maturational patterns could be caused by influences from extraneous task-specific neural processes (i.e., not specific to multisensory integration). Phenomena such as modality switch effects, which contribute significantly to multisensory gain in a bisensory detection task but not in an AV speech identification task, could prolong the perceived maturational course of multisensory processing. While this is consistent with the fact that maturation of MSEs (visual to auditory) extended well into adulthood (Fig. 7B, left), the developmental trajectory of multisensory gain was qualitatively unchanged when the contribution of MSEs was diminished by focusing our analysis on the repeat trials (Fig. S4). This, and the results of the mediation analysis, suggest that MSEs are not the sole driving factor behind the disparity in multisensory development across the two paradigms.

### Multisensory competition and sensory dominance

Human behavioral studies have demonstrated the co-occurrence of multisensory competition and facilitation using RT measures (Sinnett et al., 2008). The existence of a visual dominance in adults (i.e., the Colavita visual dominance effect; Colavita, 1974) means that directing participants to respond to either the auditory or visual component of an AV stimulus can have an inhibitive or facilitative effect on RTs, respectively (Molholm et al., 2004). However, in the same way that the race model is used as a threshold for detecting facilitative interactions, an upper statistical bound should be used to index genuine competitive interactions of this manner (Otto and Mamassian, 2012). Current neuro-computational perspectives of multisensory development suggest that competition is the default state of integration in the neonatal mammalian brain (Yu et al., 2019). Intuitively, a competition scenario would likely favor the most effective sensory modality, which in our case would be either the modality that is most dominant due to an inherent sensory bias (e.g., Colavita effect), or the preceding modality due to prior allocation of attentional resources. To gain insight into the nature of multisensory interactions in children and individuals with ASD, we tested two computational models that reflected the above scenarios of multisensory competition. We examined fits between the empirical data and model behavior that were based on a parametric weighting of the race model (facilitation) and each bias model (competition). In children with ASD aged 6–12 years, model fits suggested that the response to an AV stimulus was biased towards the previous sensory modality, potentially due to the presence of competitive interactions, whereas in their NT peers, the same models provided only marginal improvements beyond probability summation. This suggests that NT individuals develop the ability to integrate multisensory information in a facilitative manner much earlier in development than their ASD peers, who do not show evidence of facilitation until adolescence. These results are consistent with the developmental stages at which we observe the emergence of multisensory gain as indexed by violation of the race model.

Another interesting finding to emerge from our modelling analysis was that NT children aged 6–9 years appear to be biased towards the auditory modality during AV processing, but thereafter are biased towards the visual modality. The same pattern was demonstrated by a follow-up analysis that examined MSE patterns on multisensory trials. These findings are supported by previous studies that have reported an auditory dominance in infants and young children presented with AV stimuli (Lewkowicz, 1988b, a), as well as the abovementioned Colavita visual dominance effect, commonly reported in adults (Colavita, 1974). Several studies have traced the transition from an auditory to a visual dominance over the course of childhood (Robinson and Sloutsky, 2004; Nava and Pavani, 2013) and, in line with our data, suggest that this sensory reweighting occurs at around 9–10 years of age (Nava and Pavani, 2013). Sensory reweighting has also been shown to occur around 8–10 years of age for the visual and haptic modalities (Gori et al., 2008). Our modelling analysis suggests that the same trend appears to emerge in children with ASD between the ages of 6–12 years, but then reverses once more during adolescence, preferencing the auditory modality in adulthood. However, our MSE analysis suggests that a visual dominance exists initially in children with ASD, shifting to an auditory dominance in adulthood. Given the smaller sample sizes in the ASD groups, the MSE measures may be a more reliable index of sensory dominance than our modelling analysis which relies on a correlational measure. Indeed, a visual dominance has been previously reported in children with ASD (Hermelin and O’Connor, 1964; O’Connor and Hermelin, 1965), but its maturation has not yet been documented to our knowledge. In support of a visual dominance in ASD, our MSE analysis revealed that children and teenagers with ASD found it easier to switch from auditory to the visual stimuli compared to their NT peers.

### Neural basis of impaired multisensory processing in autism

Prior work by our lab suggests that the neural processes underlying multisensory integration are impaired in children with autism (Brandwein et al., 2013). Specifically, we found that EEG correlates of integration were weaker (of lower amplitude) and occurred later in the information processing hierarchy. Neural indices of integration over parieto-occipital scalp between 140–160 ms were predictive of race model violation in NT children but not in children with ASD. Using the same paradigm, we recorded intracranial electrophysiology in adults with epilepsy and demonstrated that visual stimulation influenced the phase of ongoing oscillations in auditory cortex (Mercier et al., 2015), and auditory stimulation influenced the phase of ongoing oscillations in visual cortex (Mercier et al., 2013), such that cross-sensory stimulation appears to prime ancillary sensory cortices to make them more receptive to their primary sensory input. The response to the primary sensory input (e.g., visual stimulation of visual cortex) is then enhanced for multisensory trials (Mercier et al., 2013), at least in a bisensory detection task such as the current one. Furthermore, neuro-oscillatory phase alignment across the sensorimotor network was significantly enhanced by multisensory stimulation, and was related to the speed of a response (Mercier et al., 2015). Phase resetting of ongoing neural oscillations by functionally distinct and distant neuronal ensembles is thought to be fundamental to multisensory integration (Lakatos et al., 2007; Schroeder et al., 2008; Senkowski et al., 2008; Fiebelkorn et al., 2011; Fiebelkorn et al., 2013). Impaired cross-sensory phase-resetting, as might be predicted by reduced subcortical and cortical connectivity, would likely result in impaired integrative abilities. In autism, there is evidence for such disrupted connectivity (Zeng et al., 2017; Arnold Anteraper et al., 2018), although these findings are mixed and somewhat inconclusive (Vasa et al., 2016). Nevertheless, disrupted connectivity could in turn lead to impaired cross-sensory phase-resetting and hence contribute to impaired multisensory processing in ASD. As previously mentioned, weaker cross-sensory inhibition might account for reduced MSEs in ASD (Murphy et al., 2014), possibly also due to poorer brain connectivity. Alternatively, it is possible that cross-sensory connectivity is fully intact in children with ASD, but integration of multisensory information has not yet transitioned from a state of competition, to one of facilitation, as discussed earlier (Cuppini et al., 2011; Cuppini et al., 2018). Establishing the specific neural mechanisms that underlie impaired multisensory behavior in children with ASD will likely require the use of more advanced neuroimaging techniques.

### Inhibition and prediction in modality switching

One of the unexpected findings to emerge from our MSE analysis was the reduced switch costs (auditory to visual) in ASD participants between the ages of 10–16 years. This ran contrary to a recent study (Williams et al., 2013) that reported larger switch costs in individuals with ASD of approximately the same age. Interestingly, a *post hoc* analysis of our data that focused on trials with longer ISIs (closer to that of Williams et al., 2013) led to a significant reduction in MSEs in NT individuals and only a slight reduction in individuals with ASD. This modification revealed no group differences, suggesting an interaction between group and ISI. A possible explanation for this interaction points to the so-called “trace theory” which originates from research on MSEs in individuals with schizophrenia (Zubin, 1975). This theory suggests that sensory information leaves traces of residual activity in different neuronal populations, facilitating the processing of subsequent stimuli of the same sensory modality and inhibiting the processing of stimuli of other modalities. Zubin (1975) predicted that these traces attenuate over time but persist longer in individuals with schizophrenia. If such an inhibitory cross-sensory mechanism were weaker in individuals with ASD, but persisted longer over time, it would explain the interaction that we observe here and the findings of Williams et al. (2013). Evidence in support of this theory comes from a recent study that demonstrated that individuals with ASD weight recent stimuli less heavily than NT individuals and that their perception is dominated by longer-term statistics (Lieder et al., 2019). While reduced cross-sensory inhibition would undoubtedly facilitate processing of subsequent inputs in other sensory systems and thus lead to lower MSEs in ASD, it would also likely result in greater susceptibility to distraction by task/sensory-irrelevant information. This is consistent with our modelling analysis that revealed a higher (more positive) channel dependency in ASD, suggesting they distribute attentional resources more evenly across sensory inputs. Neurophysiological evidence for this comes from previous work by our lab that demonstrated increased susceptibility to distraction by task-irrelevant stimuli in children with ASD (Murphy et al., 2014). This behavioral deficit was accompanied by a reduced neural suppression of sensory-irrelevant information, as indexed by EEG recordings of alpha-band oscillatory activity. Thus, behavioral and neurophysiological accounts of multisensory attention in ASD are consistent with a reduction in MSE.

Alternatively, reduced MSEs in ASD could be explained by differences in the ability to make predictions about the sensory environment. While individuals with ASD have been shown to utilize longer-term statistics to make predictions about their sensory environment (Lieder et al., 2019), other work suggests that they tend to overestimate the volatility of their environment at the expense of learning to build stable predictions (Lawson et al., 2017). In the current study, stimuli were presented in a random order with equal probability, meaning there was a 66.6% chance of the same unisensory input occurring on the next trial (including the AV condition). Based on these statistics, it is more efficient to predict the reoccurrence of the same signal (or part of it) on the next trial and to direct attention therein. If these statistics are not being actively used to build predictions about the modality of an upcoming stimulus, as may be the case in ASD, then the participant may be less likely to prepare for it and thus less averse to switching sensory modality. This fits well with the notion that individuals with autism rely more on bottom-up than top-down processing (Maekawa et al., 2011).

### Multisensory integration or modality switch effects?

It is well established that MSEs systematically contribute to multisensory facilitation in a bisensory detection task (Gondan et al., 2004; Van der Stoep et al., 2015a; Shaw et al., 2019). To determine the role of MSEs, we performed separate tests of the race model using switch and repeat trials. While we found that multisensory gain was much greater on switch trials than on repeat trials, there was still evidence of race model violation on repeat trials. However, it is important to consider that in the context of a mixed block design, responses on repeat trials are likely subject to residual switch effects from earlier trials (*n*-2, *n*-3, etc.). Furthermore, if we consider the impact that switching modality has on RTs, a mixed block design could be said to violate the assumption of context invariance. While it is unlikely that it would present the opportunity to change strategy from trial to trial in a top-down manner, it is conceivable that the continuously changing context (from switch to repeat conditions) could invoke disparate processing mechanisms in a bottom-up manner (for a detailed discussion, see Shaw et al., 2019). We also measured the correlation between multisensory gain and MSEs on unisensory and multisensory trials, partialling out the effects of age. There was a strong positive correlation for unisensory (but not multisensory) stimuli, as would be expected if MSEs were to impact multisensory gain systematically. This was followed up with a mediation analysis to determine whether MSEs mediated the observed relationship between age and multisensory gain. This analysis indicated only partial mediation, suggesting that neural processes other than MSEs (e.g., cross-sensory interactions) were contributing to the observed multisensory gain. Differences in the developmental trajectories of MSEs and multisensory gain lend further support to the argument that MSEs are not the sole contributor to the RSE (Gondan et al., 2004).

Another way to examine the contribution of MSEs is to remove the presence of switch trials by using a blocked design. In another study by our lab (Shaw et al., 2019), we demonstrated that RTs to simple AV stimuli do not violate the race model when the three conditions are presented in entirely separate blocks. Comparing the median RTs between blocked and mixed conditions revealed a slowing of the unisensory but not the multisensory RTs in the mixed condition that could be largely accounted for by increased RTs on switch trials. In another study that employed a blocked design, Otto and Mamassian (2012) reported evidence of race model violation, but importantly, presented AV stimuli in background noise which are more likely to recruit integrative mechanisms during bisensory detection (Wallace et al., 1996; Senkowski et al., 2011; Stevenson et al., 2012; Crosse et al., 2016). However, it is important to consider the theoretical implications of employing a blocked design. The race model test relies on the assumption of context invariance, whereby the stimulus conditions are presented in an intermixed and unpredictable fashion (Miller, 1982; Gondan and Minakata, 2016; Miller, 2016). By randomly interleaving the conditions, the participant does not know which condition to expect and presumably processes, say, an auditory signal in the same way under unisensory and multisensory conditions. Thus, violation of the race model is assumed to be caused by putatively multisensory interactions as opposed to differences in processing strategies. In contrast, when unisensory and multisensory stimuli are presented in separate blocks, there may be opportunity to employ different processing strategies in order to optimize performance. Hence, it is inherently difficult to disentangle contributions of switching and integration when examining the RSE. Violation of the race model likely involves an interplay between integrative and switching processes that carry different weights in different contexts (mixed versus blocked presentations) and under different stimulus conditions (clean versus noisy).

## Conclusions

We can draw several conclusions from the present study. 1) When assessed using the race model test, multisensory processing in individuals with ASD “normalizes” by the mid-twenties. 2) In younger children with ASD, multisensory competition appears to reflect the default mode of integration, and may take longer to transition to facilitation in individuals with ASD. 3) Differences in both MSEs and developmental patterns in sensory dominance indicate fundamental alterations in how the nervous system of children with ASD responds to even the simplest of multisensory environments. 4) Greater channel dependency in ASD suggests a more even distribution of attention across sensory inputs, possibly due to reduced cross-sensory inhibition or a reluctance to exploit short-term statistics. The current findings also make clear that there is significant work ahead of us before we fully understand the neural processes that contribute to multisensory development and how it differs in children with ASD. Complicating this endeavor, we must additionally account for individual variance and group differences in such patterns, and consider how these contribute to how the sensory environment is experienced throughout development. Here we set the stage for detailed characterization of these processes and their interactions, to in turn understand potential roadblocks to the typical development of multisensory processing in ASD, and some of the factors that might contribute to sensory reactivity in this group.

## Acknowledgements

The authors thank Juliana Bates, Natalie Russo, and the Human Clinical Phenotyping Core (HCP) for their careful clinical and cognitive phenotyping of our ASD cohort, Alice Brandwein who conducted the previous research for which most of these data were collected, Cristiano Cuppini for his helpful comments and discussions, Ana Francisco for her support with statistical analyses, Douwe Horsthuis, Elise Tavern and Danielle DeMaio for their technical assistance. We also extend our heartfelt gratitude to the participants and their families that have contributed their time to participate in this research. This work was supported by the National Institute of Mental Health of the National Institutes of Health (NIH) under award number RO1MH085322 (S.M. and J.J.F.), and the Eunice Kennedy Shriver National Institute of Child Health & Human Development of the NIH under award number U54HD090260, previously P30HD071593 (Rose F. Kennedy Intellectual and Developmental Disabilities Research Center).

## Author contributions

S.M. and J.J.F. designed the original experiment. M.J.C., S.M. and J.J.F. conceived of the current study. M.J.C. analyzed the data and produced all illustrations in consultation with S.M. and J.J.F. M.J.C. wrote the first substantive draft of manuscript. S.M. and J.J.F. provided editorial input to M.J.C. on multiple subsequent drafts.

