## Supplemental Figures and Tables for "Developmental Recovery of Impaired Multisensory Processing in Autism and the Cost of Switching Sensory Modality"

### Supplementary Figures

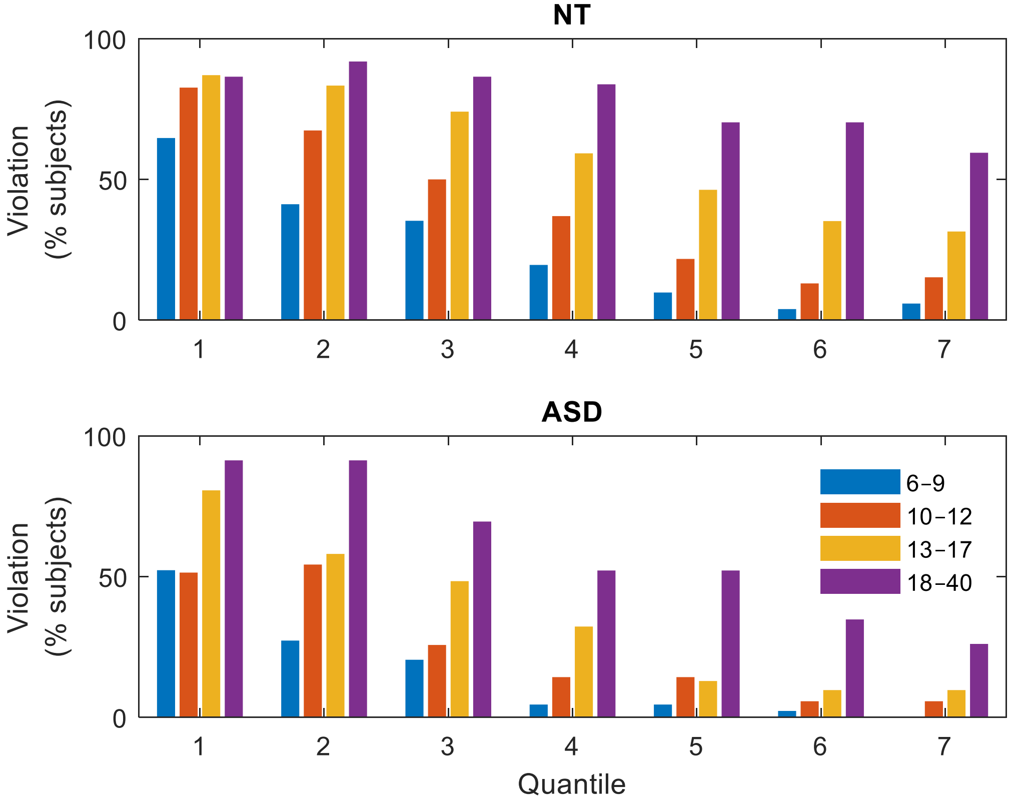


**Figure S1.** Percentage of subjects showing evidence of race model violation at each of the first 7 quantiles of the RT distribution for NT (top) and ASD (bottom) participants. Age group is indicated by color.

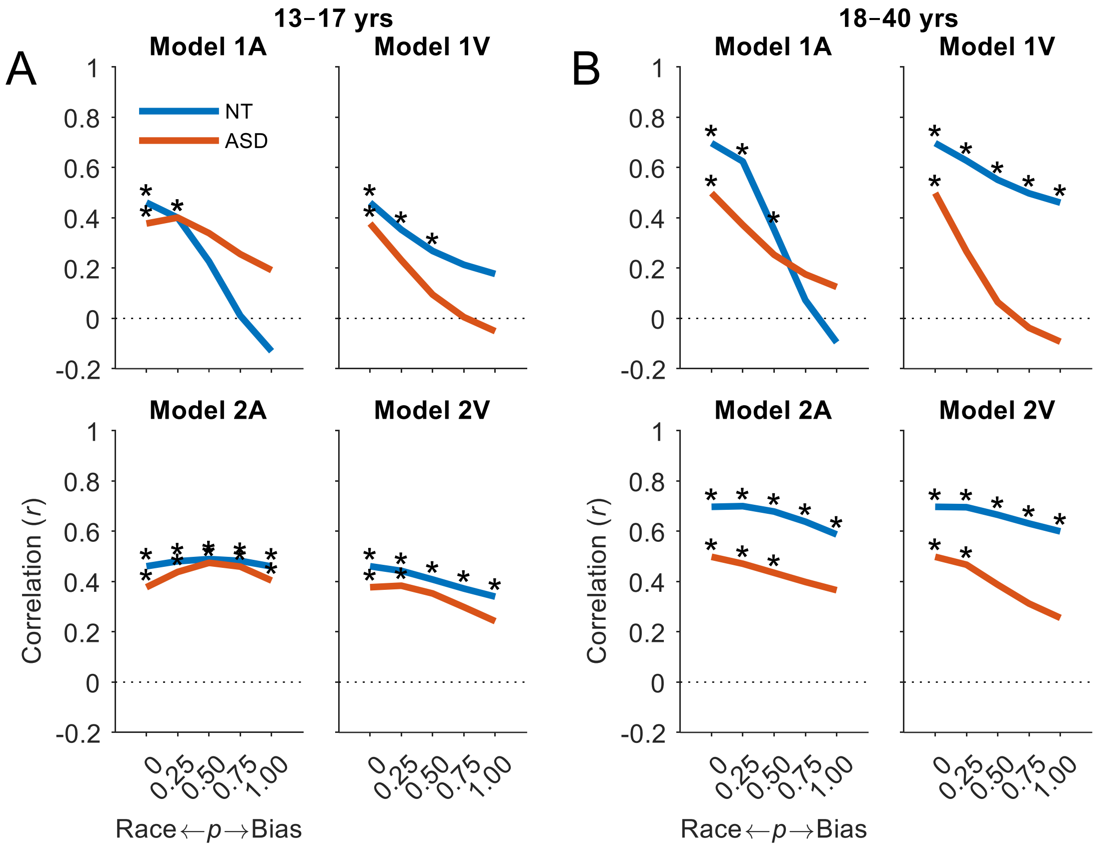


**Figure S2.** Predicting multisensory benefits in teenagers and adults. **A**, **B**, Hypothetical models of multisensory competition were tested. Model 1A was biased towards the auditory modality and Model 1V towards the visual modality. Model 2A was biased towards the preceding modality and the A modality when preceded by an AV trial, and Model 2V was biased towards the preceding modality and the V modality when preceded by an AV trial. The probability *p* of a response being triggered by a race strategy or a biased strategy was parametrically varied between 0 and 1 in increments of 0.25. The ability of each model to predict the variance in empirical benefits was assessed within each age group based on the Pearson correlation coefficient as in panel B. Data presented are the two older age groups. See main article for the two younger age groups.

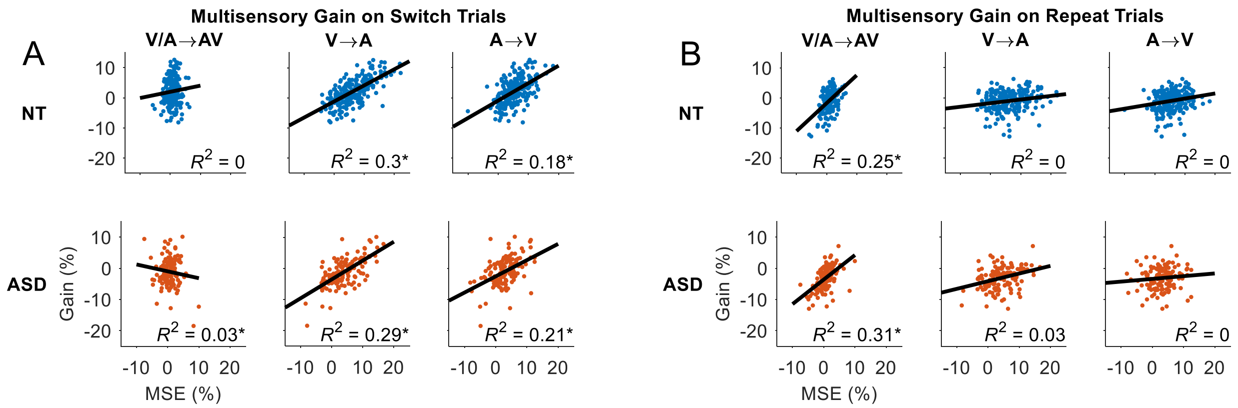


**Figure S3.** Relationship between MSEs and multisensory gain. **A**, **B**, Correlation between modality switch effects and multisensory gain on switch trials and repeat trials by condition and diagnosis (blue = NT, red = ASD). Each datapoint represents an individual participant. Explained variance ($R^2$) of the regression lines and its significance are displayed in the bottom right corner of each panel (*$p$ < 0.05; partial correlation controlling for age).

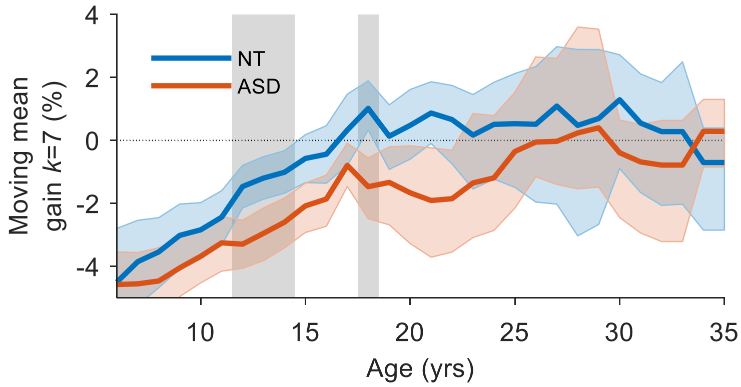


**Figure S4.** Developmental trajectory of multisnesory gain using only repeat trials. Mean multisensory gain calculated with a moving window *k* of 7 years in increments of 1 year from 6–35 years for NT (blue trace) and ASD (red trace) participants. Colored error bounds indicate 95% CIs (bootstrapped). Gray shaded regions indicate significant group differences ($p$ < 0.05, two-tailed permutation tests, FDR corrected).

### Supplementary Tables

**Table S1.** Test statistics comparing CDFs of multisensory RTs with Miller's bound. Values shown indicate effect sizes (Cohen's *d* corrected for sample size) and 95% CIs (bootstrapped) in brackets. Asterisks indicate significant race model violation ($p < 0.05$, right-tailed permutation tests, $t_{max}$ corrected).

| | NT | | | | ASD | | | |
|---|---|---|---|---|---|---|---|---|
| Q | 6–9 | 10–12 | 13–17 | 18–40 | 6–9 | 10–12 | 13–17 | 18–40 |
| 1 | 0.15[0.0,0.3]* | 0.40[0.2,0.7]* | 0.51[0.3,0.9]* | 0.83[0.7,1.1]* | 0.07[-0.1,0.3] | 0.03[-0.2,0.3] | 0.25[0.0,0.6]* | 0.60[0.3,1.1]* |
| 2 | -0.23[-0.4,-0.1] | 0.02[-0.1,0.2] | 0.23[0.1,0.5]* | 0.64[0.5,0.9]* | -0.33[-0.5,-0.2] | -0.23[-0.4,-0.1] | -0.12[-0.3,0.1] | 0.22[0.0,0.5] |
| 3 | -0.55[-0.8,-0.4] | -0.34[-0.5,-0.2] | 0.01[-0.1,0.2] | 0.33[0.2,0.5]* | -0.65[-0.9,-0.5] | -0.51[-0.8,-0.3] | -0.56[-0.9,-0.3] | -0.10[-0.3,0.1] |
| 4 | -0.89[-1.4,-0.6] | -0.67[-1.0,-0.5] | -0.28[-0.5,-0.1] | -0.01[-0.2,0.1] | -1.05[-1.6,-0.8] | -0.90[-1.5,-0.6] | -1.21[-2.2,-0.8] | -0.24[-0.6,-0.0] |
| 5 | -1.25[-2.1,-0.9] | -1.07[-1.7,-0.8] | -0.56[-0.9,-0.4] | -0.44[-0.7,-0.3] | -1.36[-2.5,-0.9] | -1.20[-2.1,-0.9] | -1.92[-3.4,-1.3] | -0.35[-0.8,-0.1] |
| 6 | -1.84[-2.8,-1.4] | -1.51[-2.2,-1.2] | -0.79[-1.4,-0.5] | -0.90[-1.3,-0.6] | -1.68[-3.0,-1.2] | -1.58[-2.9,-1.1] | -2.62[-3.8,-2.0] | -0.47[-1.1,-0.3] |
| 7 | -2.11[-2.8,-1.7] | -1.68[-2.3,-1.4] | -1.08[-1.8,-0.7] | -1.33[-2.0,-0.9] | -2.09[-3.1,-1.5] | -1.59[-2.7,-1.2] | -2.65[-3.5,-2.1] | -0.70[-1.4,-0.5] |

**Table S2.** Test statistics comparing CDFs of multisensory RTs with the race model using only repeat trials. Values shown indicate effect sizes (Cohen's *d* corrected for sample size) and 95% CIs (bootstrapped) in brackets. Asterisks indicate significant race model violation ($p < 0.05$, right-tailed permutation tests, $t_{max}$ corrected).

| | NT | | | | ASD | | | |
|---|---|---|---|---|---|---|---|---|
| Q | 6–9 | 10–12 | 13–17 | 18–40 | 6–9 | 10–12 | 13–17 | 18–40 |
| 1 | 0.28[0.1,0.5]* | 0.48[0.3,0.8]* | 0.48[0.3,0.7]* | 0.73[0.5,1.0]* | 0.10[-0.1,0.3] | -0.02[-0.3,0.3] | 0.17[-0.0,0.5] | 0.78[0.5,1.2]* |
| 2 | -0.13[-0.3,0.0] | 0.14[-0.0,0.3] | 0.27[0.1,0.4]* | 0.50[0.4,0.7]* | -0.24[-0.4,-0.1] | -0.09[-0.3,0.1] | -0.06[-0.3,0.1] | 0.24[0.0,0.5] |
| 3 | -0.27[-0.4,-0.1] | -0.09[-0.2,0.1] | 0.11[-0.0,0.2] | 0.38[0.2,0.5]* | -0.44[-0.6,-0.3] | -0.29[-0.5,-0.1] | -0.15[-0.4,0.1] | 0.02[-0.2,0.2] |
| 4 | -0.45[-0.7,-0.3] | -0.25[-0.4,-0.1] | -0.02[-0.2,0.1] | 0.20[0.1,0.3]* | -0.60[-0.9,-0.4] | -0.44[-0.8,-0.3] | -0.36[-0.7,-0.2] | -0.10[-0.3,0.1] |
| 5 | -0.65[-1.0,-0.5] | -0.42[-0.6,-0.3] | -0.17[-0.3,-0.0] | -0.01[-0.2,0.1] | -0.74[-1.2,-0.5] | -0.60[-1.0,-0.4] | -0.60[-1.0,-0.3] | -0.19[-0.5,0.0] |
| 6 | -0.88[-1.3,-0.7] | -0.57[-0.8,-0.4] | -0.24[-0.4,-0.1] | -0.13[-0.3,0.0] | -0.99[-1.5,-0.7] | -0.72[-1.1,-0.5] | -0.86[-1.4,-0.5] | -0.23[-0.6,-0.1] |
| 7 | -1.01[-1.4,-0.8] | -0.69[-1.0,-0.5] | -0.38[-0.7,-0.2] | -0.28[-0.5,-0.1] | -1.11[-1.7,-0.7] | -0.73[-1.1,-0.5] | -0.98[-1.5,-0.6] | -0.35[-0.7,-0.2] |

**Table S3.** Test statistics comparing CDFs of multisensory RTs with Miller's bound using only repeat trials. Values shown indicate effect sizes (Cohen's *d* corrected for sample size) and 95% CIs (bootstrapped) in brackets. Asterisks indicate significant race model violation ($p < 0.05$, right-tailed permutation tests, $t_{max}$ corrected).

| | NT | | | | ASD | | | |
|---|---|---|---|---|---|---|---|---|
| Q | 6–9 | 10–12 | 13–17 | 18–40 | 6–9 | 10–12 | 13–17 | 18–40 |
| 1 | 0.23[0.1,0.4]* | 0.44[0.2,0.8]* | 0.44[0.2,0.7]* | 0.73[0.5,1.0]* | 0.06[-0.1,0.2] | -0.07[-0.4,0.3] | 0.13[-0.1,0.4] | 0.76[0.5,1.2]* |
| 2 | -0.31[-0.5,-0.1] | -0.07[-0.2,0.1] | 0.10[-0.1,0.3] | 0.37[0.2,0.6]* | -0.42[-0.6,-0.3] | -0.25[-0.5,-0.1] | -0.30[-0.6,-0.1] | 0.06[-0.2,0.3] |
| 3 | -0.61[-0.9,-0.4] | -0.49[-0.7,-0.3] | -0.22[-0.4,-0.1] | 0.07[-0.1,0.2] | -0.81[-1.1,-0.6] | -0.63[-1.0,-0.4] | -0.66[-1.1,-0.4] | -0.28[-0.6,-0.0] |
| 4 | -0.97[-1.5,-0.7] | -0.83[-1.2,-0.6] | -0.47[-0.7,-0.3] | -0.33[-0.5,-0.2] | -1.15[-1.7,-0.8] | -0.99[-1.7,-0.7] | -1.31[-2.3,-0.9] | -0.46[-0.9,-0.2] |
| 5 | -1.31[-2.1,-1.0] | -1.32[-2.0,-1.0] | -0.72[-1.1,-0.5] | -0.89[-1.2,-0.6] | -1.40[-2.5,-0.9] | -1.29[-2.2,-0.9] | -2.04[-3.3,-1.4] | -0.57[-1.2,-0.3] |
| 6 | -1.78[-2.6,-1.3] | -1.68[-2.2,-1.4] | -0.85[-1.5,-0.5] | -1.36[-2.0,-1.0] | -1.75[-2.9,-1.2] | -1.69[-2.5,-1.3] | -2.57[-3.4,-2.1] | -0.62[-1.4,-0.4] |
| 7 | -1.85[-2.4,-1.5] | -1.69[-2.1,-1.4] | -1.11[-1.8,-0.7] | -1.53[-2.1,-1.1] | -1.88[-2.8,-1.3] | -1.50[-2.1,-1.2] | -2.09[-2.8,-1.7] | -1.05[-1.8,-0.8] |
